# Diploid genome assembly of human fibroblast cell lines enables clone specific variant calling, improved read mapping and accurate phasing

**DOI:** 10.1101/2025.04.15.648829

**Authors:** T. Rhyker Ranallo-Benavidez, Yue Hao, Emilia Volpe, Maryam Jehangir, Noelle Fukushima, Ogechukwu Mbegbu, Rebecca Reiman, Jessica Molnar, Danyael Murphy, Dorothy Marie Paredes, Shukmei Wong, Kara Karaniuk, Stephanie Buchholtz, Jonathan Keats, Mitchell J. Machiela, Mikhail Kolmogorov, Benedict Paten, Simona Giunta, Floris P. Barthel

**Author notes:** These authors contributed equally to this work (T. Rhyker Ranallo-Benavidez, Yue Hao).

## Abstract

Human cell lines are fundamental tools in biomedical research and are widely used in disease modeling, drug development, and many other domains. Here, we present chromosome-level, phased diploid genome assemblies of two popular human cell lines: the BJ foreskin fibroblast line and the IMR-90 fetal lung fibroblast line. Our high-quality assemblies, generated using long-read and Hi-C sequencing data, reveal substantial structural variation, including more than 50,000 insertions, deletions, duplications, and inversions compared to the recent T2T-CHM13v2.0 reference. Our assemblies provide detailed maps of genetic variation, enabling more accurate variant calling and the ability to phase reads when using newly generated or historical sequencing data on these cell lines or their derivatives. All assemblies and associated data have been made available as a resource for the research community. We envision that diploid genome assembly will become a cornerstone approach for personalized medicine in the near future.

## Introduction

Human cell lines are essential tools driving innovation in biomedical research. Their adaptation to *in vitro* culture conditions allows them to be used as robust model systems to study response to diverse experimental conditions, understand fundamental cellular processes, and elucidate disease mechanisms^1^. Two widely utilized cell lines are the finite BJ and IMR-90 human fibroblast lines, which exhibit a stable diploid karyotype over many population doublings but enter permanent growth arrest (senescence) after a fixed number of passages^2,3^. The BJ cell line (46,XY), derived from neonatal foreskin tissue, has been used to study telomere shortening^4^, the derivation of stem cell lines^5^, chromatin accessibility^6^, and the premature aging disease Hutchinson–Gilford progeria syndrome^7^. The IMR-90 cell line (46,XX), derived from fetal lung tissue, has been used to study cell motility^8^, viral susceptibility^3^, senescence^9^, and chromosome instability in cancer progression^10^.

Despite having been used in research for several decades, the complete genetic background of these foundational cell lines remains largely unexplored at a haplotype-resolved level. The accurate interpretation of biological experimentation in cell lines requires a comprehensive understanding of their genetic context. While the GRCh38^11^ and, recently, T2T-CHM13v2.0^12^ reference genomes provide a valuable framework, they do not capture the extensive genomic diversity present across individuals and, importantly, within laboratory cell lines. Recognizing this limitation, the Human Pangenome Reference Consortium (HPRC) aims to create a more complete and diverse human pangenome reference by assembling high-quality, phased genome sequences from hundreds of individuals representing global genetic diversity^13^. The HPRC has made significant strides in generating highly contiguous, telomere-to-telomere assemblies, revealing previously uncharacterized structural variation and improving the representation of complex genomic regions^14^.

The current study complements and extends the critical work of the HPRC by focusing on the *de novo* assembly of diploid genomes of the BJ and IMR-90 cell lines. While the HPRC focuses on assembling genomes from diverse human individuals, our work addresses a distinct, yet equally important, need: characterizing the specific genomic landscapes of widely used *in vitro* model systems. Cell lines, through their derivation and propagation over time, are thought to accumulate substantial genomic alterations, including single nucleotide variants (SNVs), insertions and deletions (indels), and, crucially, large-scale structural variants (SVs) that can significantly impact cellular phenotypes and experimental outcomes^15^. These alterations are often unique to a specific cell line and are not represented in population-based pangenome references.

Here, we present publicly available chromosome-level assemblies of the BJ and IMR-90 cell line. Crucially, the assemblies are diploid, meaning that both parental haplotypes are resolved for each cell line. This haplotype resolution is essential for accurately representing and phasing genetic variants, identifying allele-specific gene expression, and understanding the functional consequences of structural variation^16^. Many previous genomic studies have relied on mapping to the reference genome, which can lead to biases and incomplete or inaccurate variant calling, particularly in regions of high divergence or structural complexity^17^. Our *de novo* diploid assemblies overcome these limitations, enabling accurate reanalysis of previously generated sequencing data for these cell lines. By providing chromosome-level, phased genomes for BJ and IMR-90, we enable a more nuanced understanding of their biology and facilitate more accurate and reproducible research using these cell lines and their derivatives.

## Results

### Long-read and chromatin conformation capture sequencing produced chromosome-level diploid genome assemblies

We first generated sequencing libraries from BJ and IMR-90 fibroblast cells (**Figure 1A, Supplementary Figure 1A**). The generated sequencing data included Oxford Nanopore Ultra Long (ONT-UL), PacBio HiFi, and Hi-C (**Table 1**). NanoPlot^18^ sequencing quality control analysis indicated that this data was of high quality (**Table 1**, **Supplementary Figure 1B**).

**Figure 1:**
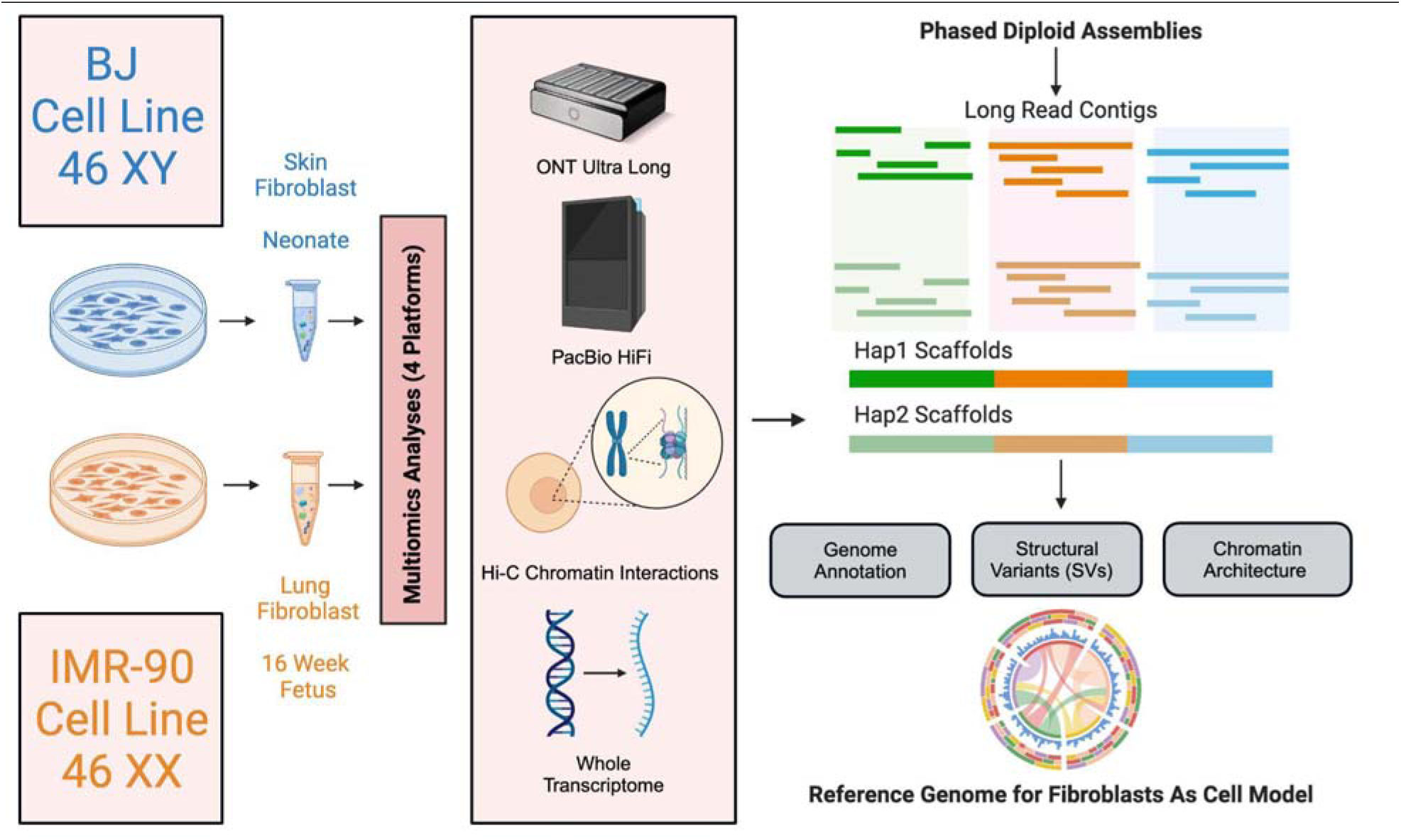
Multi-omic *de novo* diploid genome assembly workflow. Schematic showing our approach to the diploid genome assembly of two widely utilized laboratory fibroblast cell lines derived from a 46,XY and 46,XX donor. The assemblies can be used as reference genomes for studies using BJ, IMR-90, or derivative lines as model systems.

**Table 1:** Total coverage, read length N50, and median quality scores for all sequencing data. These values were calculated using NanoPlot. Total coverage was based on the T2T-CHM13v2.0 genome size of 3,117,292,070 bp. The values for the Hi-C data were calculated after trimming five bases from the 5’ end of both read 1 and read 2 as recommended by the Arima mapping pipeline.

| Sample | Data | Total Coverage | Read Length N50 | Median Quality Score |
| --- | --- | --- | --- | --- |
| BJ | ONT-UL | 25.36x | 66,433 | 20.7 |
| BJ | HiFi | 44.02x | 15,421 | 33.2 |
| BJ | Hi-C | 25.69x | 146 | 32.5 |
| IMR-90 | ONT-UL | 21.22x | 143,098 | 20.9 |
| IMR-90 | HiFi | 32.32x | 16,449 | 33.8 |
| IMR-90 | Hi-C | 26.35x | 146 | 32.7 |

As an independent, alignment-free quality metric of the long-read data, we produced k-mer spectra using GenomeScope 2.0^19^ (**Supplementary Figure 1C**). This analysis confirmed the sequencing coverages and expected heterozygosity levels for all the long-read data noted previously. Importantly, our ONT and HiFi data showed a large tail of ultra-long reads which helped the assembly of repetitive regions (**Supplementary Figure 1D**). Finally, evaluation of our Hi-C data showed a healthy distribution of long-range contacts (**Supplementary Figure 1E**)^20^.

We produced *de novo* diploid genome assemblies using Verkko 2.2.1^21^. Pre-curation assembly graph visualization and quality control suggested that our diploid assemblies were approaching the haploid T2T-CHM13v2.0 reference genome in quality (**Supplementary Figures 2-3, Supplementary Table 1**). To improve our assemblies further, we manually curated both assemblies using the Hi-C maps to correct misjoins and consolidate segments from the same chromosomes into a single scaffold (see **Methods**). This refinement process substantially improved the structural quality of the assembly, ensuring that each chromosome was accurately reconstructed as a continuous and well-defined scaffold.

Following manual curation, the BJ and IMR-90 diploid assemblies spanned 5.9 Gbp and 6.0 Gbp, respectively. Both assemblies consisted of 46 scaffolds representing the 44 autosomes and two sex chromosomes (XY for BJ and XX for IMR-90). The number of gaps were 53, 39, 41, and 29, which corresponded to less than 0.085%, 0.052%, 0.053%, and 0.056% of the genome length, respectively. To quantify the contiguity of the assemblies (see **Methods** for a description of the contiguity metrics), we calculated the auN values which represent the area under the contiguity (Nx) curve. The scaffold auN values ranged from 152.6 Mbp to 156.1 Mbp, with three out of four haplotypes exceeding the 154.6 Mbp auN value of the T2T-CHM13v2.0 reference (**Table 2**, **Figure 2A**). Using compleasm^22^ to evaluate conserved single-copy orthologous genes, we found our assemblies were over 99% complete, except for BJ hap1, which was over 96% complete (**Figure 2B**).

**Figure 2:**
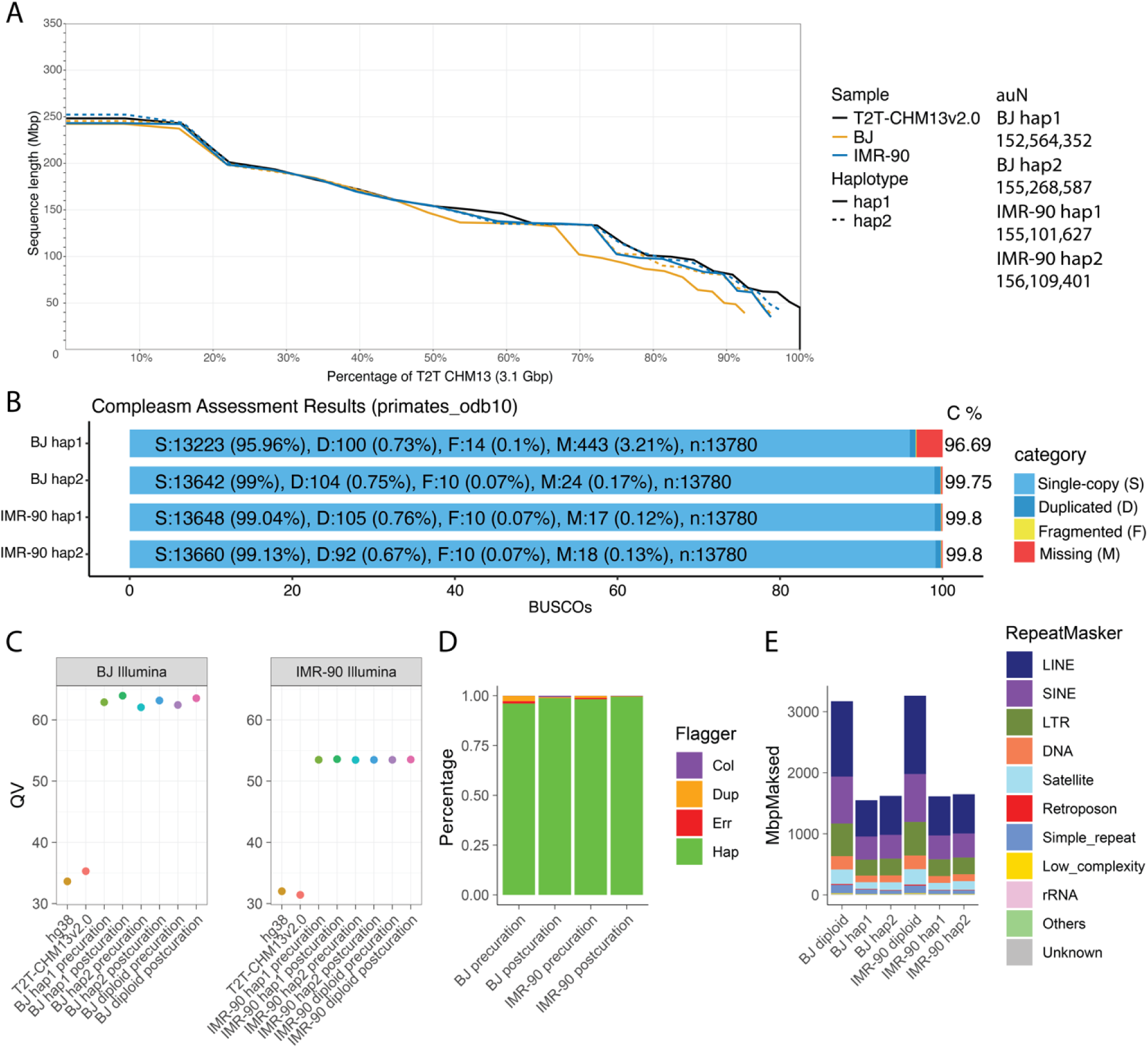
Post-curation assembly QC. **A** NGx chart showing the contiguity of the four post-curation assembly haplotypes (based on the genome size of the T2T-CHM13v2.0 reference). The NG50 value (when x=50%) is the length of the shortest sequence needed to cover at least 50% of the total reference assembly. In black is the NGx plot for the T2T-CHM13v2.0 reference. To the right of the chart are the auN metrics for the post-curation haplotypes. **B** The compleasm results showing the estimated number of single-copy (S), duplicated (D), fragmented (F), and missing (M) near-universal single-copy ortholog gene sequences in the post-curation haplotypes. **C** Merqury estimated QV of the pre- and post-curation haplotypes using publicly available Illumina data. **D** HMM-Flagger results showing the percentage of each pre- and post-curation haplotype that is categorized as correctly assembled (hap), in error (err), falsely duplicated (dup), or collapsed (col). **E** RepeatMasker results showing the number of bases in the post-curation haplotypes that are categorized into each repeat family.

**Table 2:**
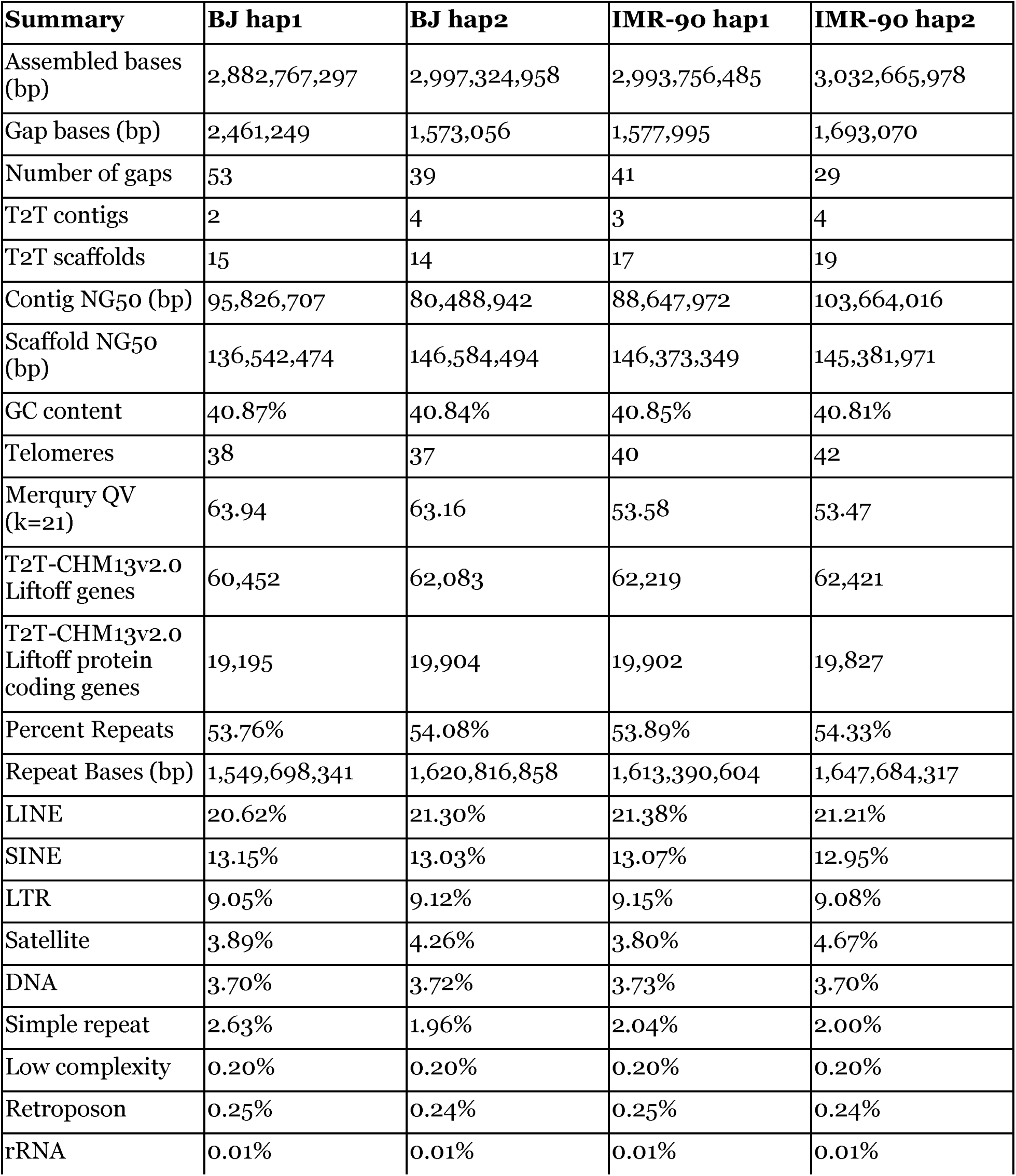
Assembly statistics for the post-curation assemblies. The assembled bases, gap bases, number of gaps, contig and scaffold NG50s, and GC content were calculated using gfastats. The QV scores were calculated using Merqury with publicly available Illumina data. The T2T-CHM13v2.0 gene annotation was lifted over to each post-curation assembly haplotype using Liftoff. The repeat content was calculated using RepeatMasker.

We next ran Merqury^23^ using publicly available BJ and IMR-90 Illumina whole genome sequencing data to determine the consensus quality values (QV), indicating the correctness of our assemblies (**Figure 2C**). For BJ and IMR-90 respectively, the QV scores were 63.53 and 53.52 and the completeness scores were 99.43% and 99.46%. In comparison, we also quantified correctness when comparing the same publicly available BJ and IMR-90 short reads to non-matching reference genomes. QV scores for GRCh38 and T2T-CHM13v2.0 ranged between 32 and 36 when using BJ or IMR-90 short-read data. These results show that the BJ and IMR-90 assemblies more accurately represent corresponding BJ and IMR-90 short-read data by more than two orders of magnitude when compared to an unmatched reference.

We then used Flagger^13^ to determine the percentage of each assembled haplotype that is categorized as correctly assembled (haploid), in error, falsely duplicated, or collapsed (**Figure 2D**). Our final BJ and IMR-90 assemblies were predicted to be over 98% and 99% correctly assembled, respectively. Final inspection of the Hi-C contact map showed characteristic diamond shapes for each of the chromosomes reflecting spatially segregated haplotypes and did not reveal any off-diagonal contacts, confirming the correct order and orientation of all scaffolds (**Supplementary Figure 4**).

We finally used RepeatMasker^24^ to locate and quantify the repeat families within each of the assembled haplotypes (**Figure 2E**). The number of bases in each category closely matched the corresponding numbers in the T2T-CHM13v2.0 reference, with LINEs comprising around 21% of each haplotype, and SINEs comprising around 13%. LTRs comprised the next most prevalent repeat family, with around 9% in each haplotype.

### Computational karyograms confirm the structural integrity of the assemblies

To evaluate the structural completeness of our assemblies and assign chromosome identifiers, we compared traditional karyotypes to our assemblies. Multiplex fluorescence *in situ* hybridization (M-FISH) karyotyping of BJ and IMR-90 cells confirmed that both lines were diploid and 46,XY and 46,XX, respectively (**Figure 3A-B**). To facilitate the comparison of traditional karyotypes and assembled genomes, we developed KaryoScope. KaryoScope annotates the k-mers in a genomic sequence based on their location in the T2T-CHM13v2.0 reference (see **Methods**).

**Figure 3:**
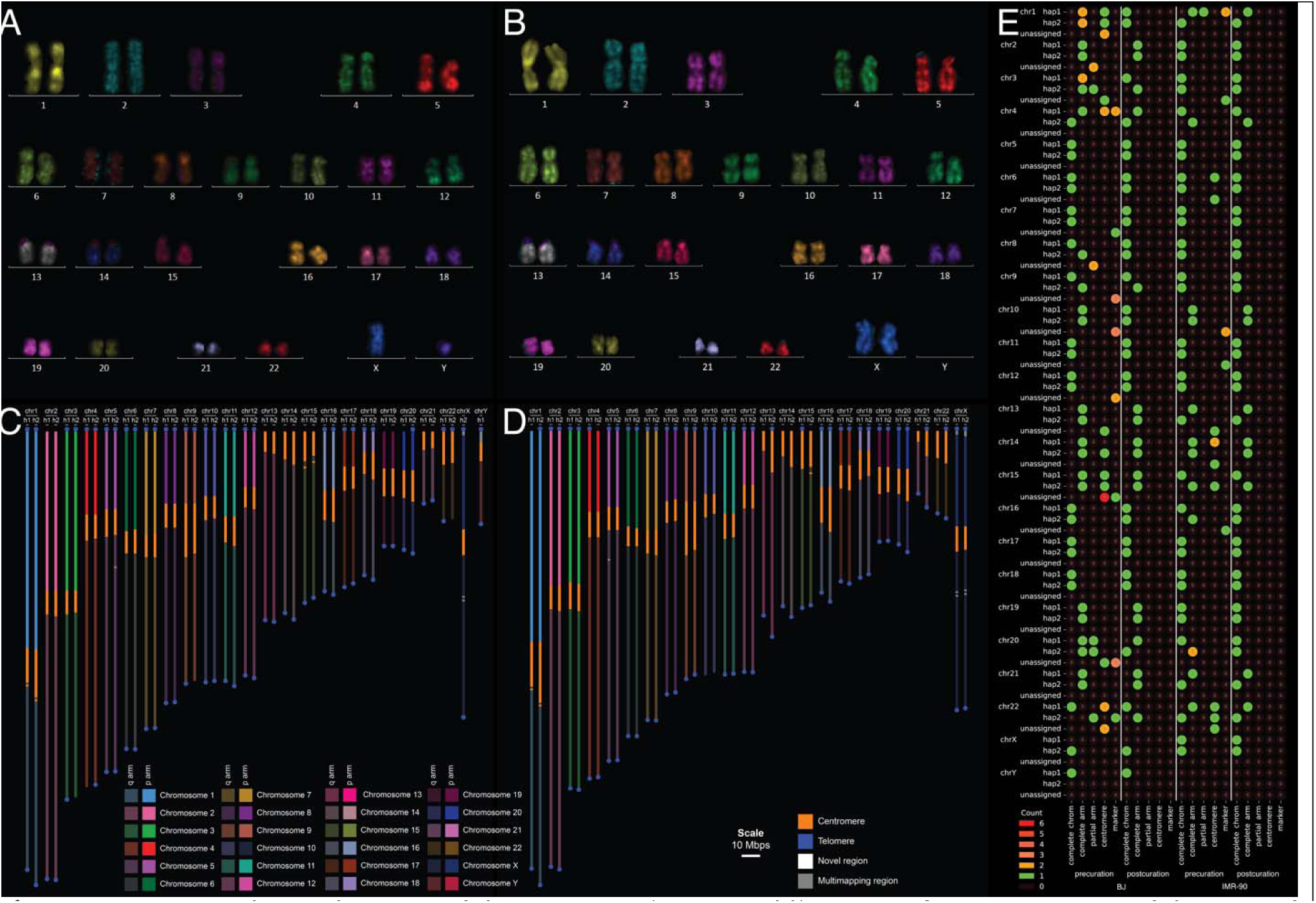
Structural completeness of the post-curation assemblies. **A and B** Karyograms of the BJ and IMR-90 cell lines, respectively, acquired using the MetaSystems Metafer Slide Scanning Platform and Ikaros module. **C and D** Computational karyograms of the BJ and IMR-90 post-curation assemblies. **E** Status of completeness for the pre- and post-curation assemblies. Each contig/scaffold of the assembly is categorized as complete chromosome, complete arm, partial arm, centromere, or marker.

We used KaryoScope to identify the chromosome arms, centromeres, and telomeres in the pre- and post-curation assemblies (**Supplementary Figure 5, Figure 3C-D**). Although pre-curation assemblies were broken at a couple centromeres, the post-curation assemblies were almost entirely telomere-to-telomere. Some chromosomes, including the acrocentric p-arms, were missing telomeres in our final assembly, but the rest of the chromosome was intact. Overall, the genome structure observed in our genome assemblies closely mirrored what we saw in the traditional karyotypes.

The computational karyograms confirmed that the final assemblies consisted of one scaffold per pseudomolecule. To quantify structural completeness, we categorized every contig or scaffold of the pre- and post-curation assemblies into one of five categories. “Complete chromosome” comprises fully telomere-to-telomere sequences. “Complete arm” comprises sequences with at least one complete arm bound by a centromere and a telomere. The categories “partial arm”, “centromere” and “marker chromosome” were reserved for sequences that did not meet these stricter criteria. For the post-curation assemblies, all scaffolds were either classified as “complete chromosome” when complete or “partial chromosome” if they were missing one of the telomeres (**Figure 3E, Supplementary Table 2**).

### De novo assemblies resolve complex centromere and telomere regions

Centromeres and telomeres have routinely been omitted from reference genomes due to their repetitive nature and the complexities associated with their assembly. This has all changed with the efforts of the T2T and HPRC consortia^25,26^. To validate the quality and correctness of our centromere assemblies, we analyzed centromeric sequence homology using StainedGlass^27^ for both haplotypes of BJ and IMR-90 (chromosomes 5 and 11 shown in **Figure 4A-B**). Our Flagger results showed that 21 and 35 of the characteristic higher order repeat (HOR) arrays of the centromeres were correctly assembled in BJ and IMR-90, respectively. From the StainedGlass plots, we found that the structures of the functionally active HOR arrays were conserved between the two cell lines, and that the HOR units and monomers were similar to the centromere structures seen in the T2T-CHM13v2.0 reference genome^25^.

**Figure 4:**
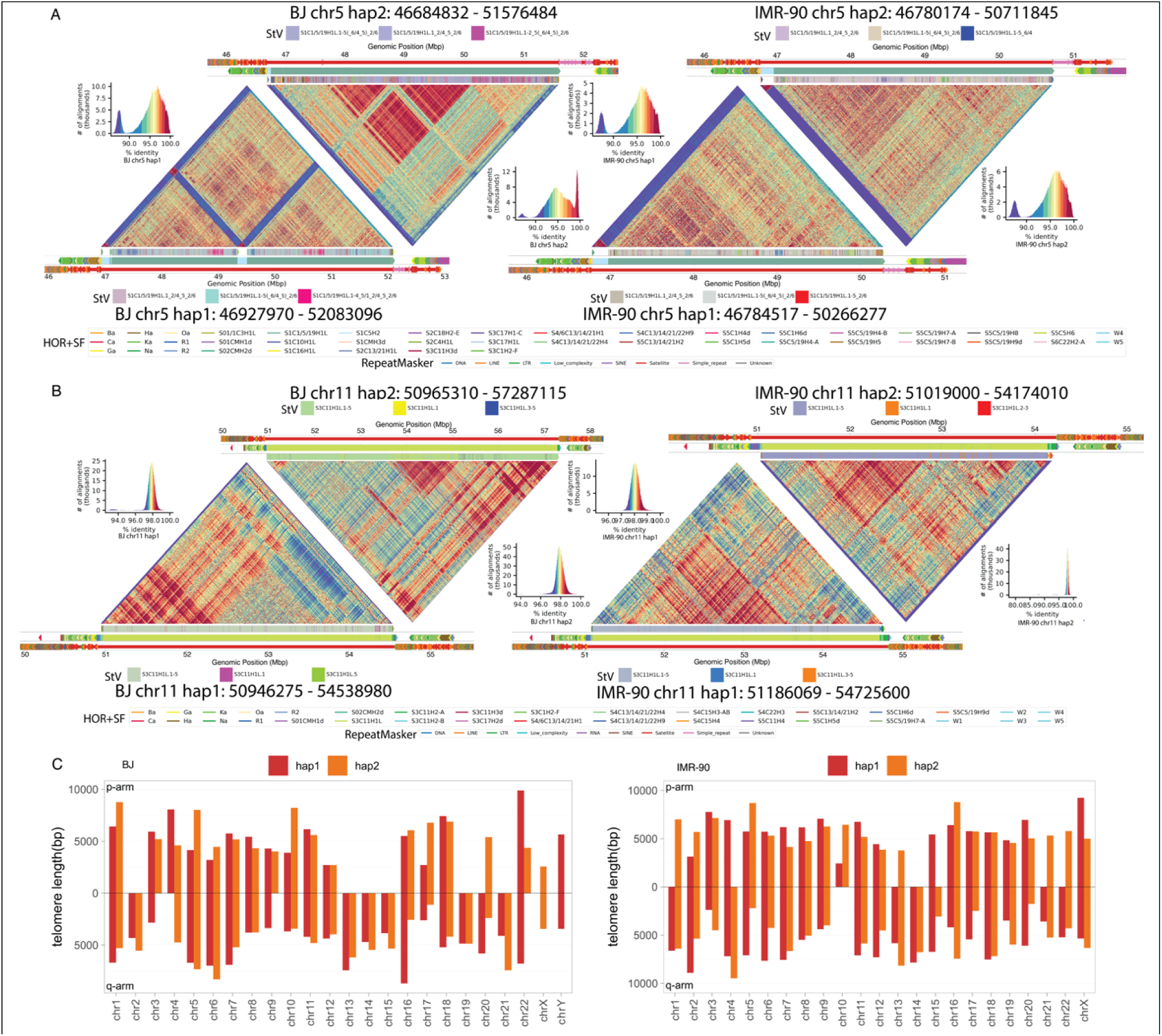
Centromeres and telomeres in the post-curation assemblies. **A and B** StainedGlass plots of BJ and IMR-90 α satellite higher-order repeats (HORs) within the chr5 and chr11 centromeres. The colors in each heatmap are based on sequence alignment percent identity (shown in the histograms), with red being the highest level of sequence similarity. The innermost annotation track consists of the live HORs only. Each color represents a different StV variant of the HOR monomers. The top three most abundant variants within each centromere are labeled. The central annotation track is an extended region including 1Mbp upstream and downstream of the live HORs. Each color represents a different HOR or other repeat superfamily (HOR+SF). The outermost annotation track consists of the repeat classes from RepeatMasker. **C** Bar plots of arm-specific telomere lengths for BJ (left) and IMR-90 (right).

The immediate flanking satellite sequence upstream and downstream of the active HOR arrays was also conserved within each chromosome and compared to T2T-CHM13v2.0. These results indicate that some regions within human centromeres are highly conserved. This is not surprising as any disruptions would have detrimental consequences for cell division. Within the active HOR arrays of each centromere, we annotated the different variants of alpha satellites and found that the two haplotypes in BJ showed more differences in the arrangement of HOR units than those in IMR-90. The more divergent centromere haplotypes between the two parents of BJ might suggest different ancestral history.

Like centromeres, telomeres and subtelomeres have long been excluded from human reference genomes. Our assemblies included *n* = 75 telomeres in BJ and *n* = 82 telomeres in IMR-90 out of *n* = 92 chromosome arms (**Figure 4C**). Many of the missing telomeres were located on the acrocentric arms, with *n* = 8 missing acrocentric telomeres in BJ and *n* = 6 missing acrocentric telomeres in IMR-90 out of *n* = 10 total acrocentric telomeres.

We found no difference in telomere length comparing the two haplotypes in each cell line. IMR-90 telomeres were slightly longer than BJ, possibly suggesting that our BJ cells spent less time in culture prior to our handling (**Supplementary Figure 6A**). Compared to the HPRC samples, we found that BJ and IMR-90 telomeres were significantly shorter, most likely because BJ and IMR-90 cell lines spent extensive time in culture relative to low-passage HPRC cell lines. This hypothesis is supported by the fact that the HPRC telomeres were shorter in later project years (**Supplementary Figure 6B**). Early HPRC samples were held strictly to minimal passaging whereas this constraint was relaxed in later project years (personal communication).

### Structural comparisons of the assemblies reveal large scale variations within and between cell lines

Multi-sample synteny analysis of the 22 autosomes across our assemblies and the T2T-CHM13v2.0 reference^28^ revealed a high degree of collinearity of the two *de novo* diploid assemblies (**Figure 5A**). Both BJ and IMR-90 contained a large haplotype-specific inversion on chromosome 8 (hap1) when compared to the T2T-CHM13v2.0 reference. We confirmed the existence of both breakpoints of the inversion in both cell lines (IMR-90 shown in **Supplementary Figure 7**). The inversion on chromosome 8 in human was previously reported to be associated with miscarriages when one parent carries such structural variation^29^. To check for large-scale haplotype-specific structural variants within each cell line, we aligned both haplotypes to each other and visualized the alignments using SVbyEye^30^ (**Figure 5B-C**). Although the inversions on chromosome 8 are clearly visible, no other obvious large-scale haplotype-specific variation could be observed.

**Figure 5:**
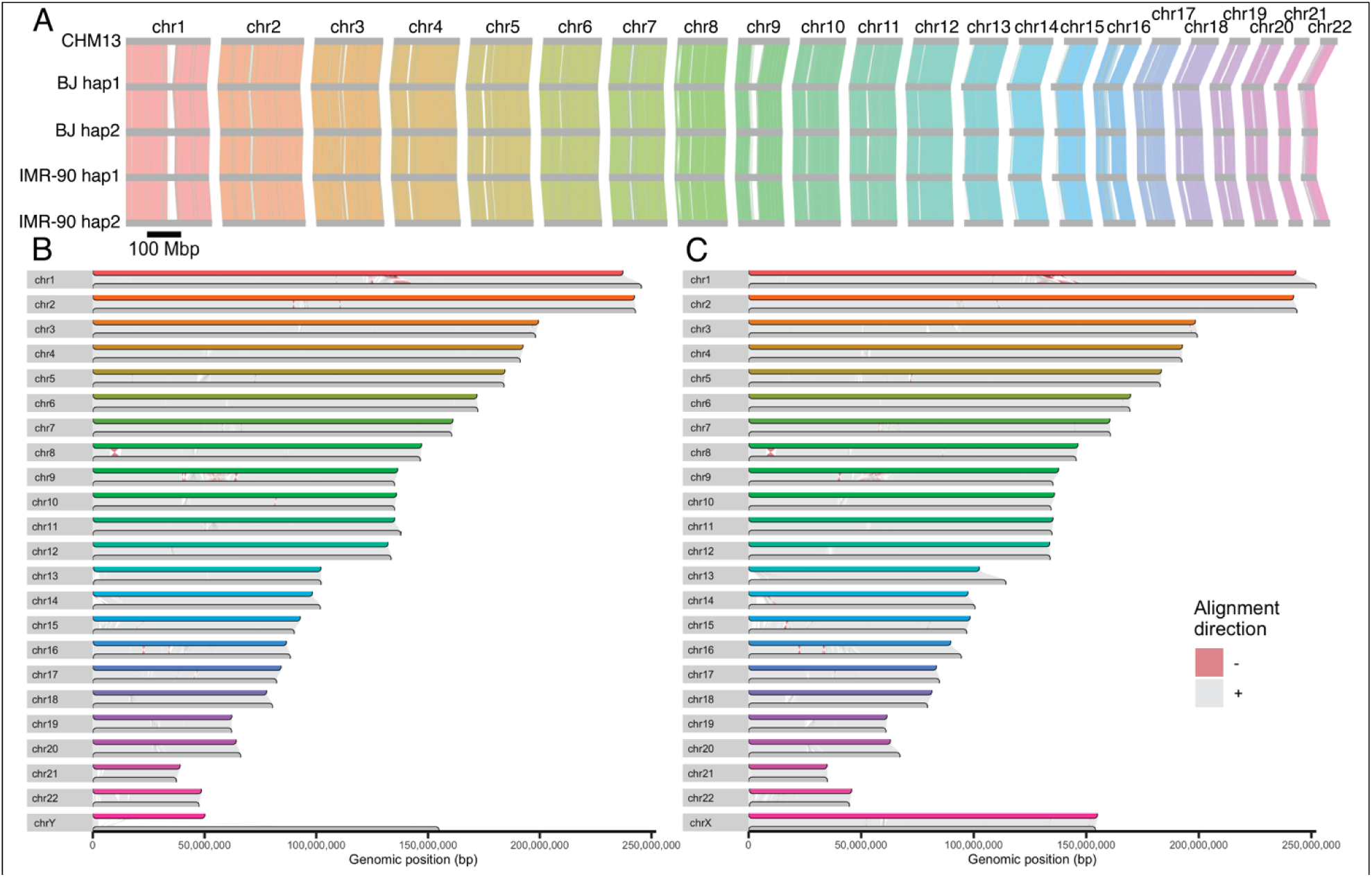
Post-curation assembly synteny plots. **A** Multi-sample synteny between the T2T-CHM13v2.0 reference and the four post-curation assembly haplotypes generated by ntSynt. Sex chromosomes not shown. Conserved blocks are indicated by colored links between haplotypes. **B and C** Haplotype1 versus haplotype2 synteny plots for BJ and IMR-90 post-curation assemblies, respectively, generated by SVbyEye.

To systematically call variants directly from our assemblies, we used SyRI^31^ to call SNVs, indels, and SVs within each of the four assembled haplotypes compared to the haploid T2T-CHM13v2.0 reference (**Figure 6**). After merging small variants (SNVs and indels) from four haplotypes, we found *n* = 7,786,003 SNVs and *n* = 1,486,112 indels. Amongst all small variants found in BJ (*n* = 5,611,994) and IMR-90 (*n* = 6,842,223) (**Figure 6A**), *n* = 4,003,178 (71.3%) and *n* = 5,111,723 (74.7%) were heterozygous, respectively. A total of *n* = 3,182,102 small variants (56.7% of BJ and 46.5% of IMR-90) were shared between cell lines. Conversely, *n* = 2,429,892 (43.3%) of small variants found in BJ were specific to BJ whereas *n* = 3,660,121 (53.5%) of those found in IMR-90 were specific to IMR-90. To confirm whether the BJ and IMR-90 SNVS and indels were novel or present in the general population, we compared them to gnomAD^32^ (**Figure 6B**). We found that 71.7% of SNVs in BJ, 90.1% of indels in BJ, 65.2% of SNVs in IMR-90, and 87% of indels in IMR-90 were previously noted in the gnomAD database.

**Figure 6:**
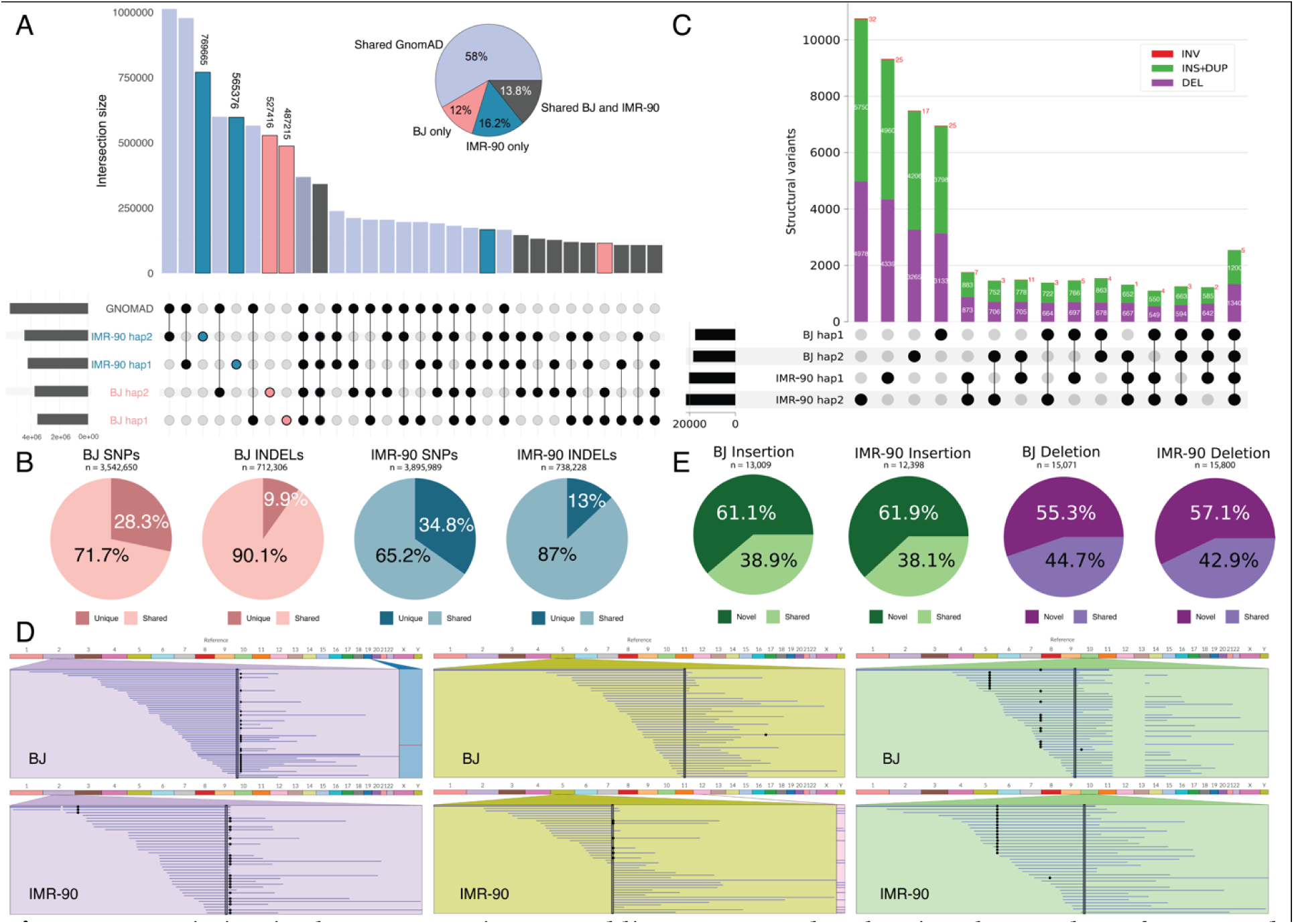
Variation in the post-curation assemblies. A UpSet plot showing the number of SNVs and indels across the four post-curation haplotypes and gnomAD. The pie chart shows the proportion of BJ and IMR-90 variants that are shared with gnomAD. **B** The percentages of BJ and IMR-90 SNVs and indels that are cell-line specific or are shared within gnomAD. **C** UpSet plot showing the number of SVs by type across the four post-curation haplotypes. **D** Three example structural variants: ribbon plot of a 4,636 bp insertion on chromosome 2 shared by all four haplotypes; ribbon plot of a 4,459 bp insertion on chromosome 5 present only in IMR-90; ribbon plot of a 4,342 bp deletion on chromosome 10 present only in BJ. **E** The percentages of BJ and IMR-90 insertion and deletion SVs that are cell-line specific or are shared within the pangenome.

We next sought to characterize large variants (SVs) found in our assemblies. After merging the four sets of SVs using Jasmine^33^, there were *n* = 51,105 SVs, including *n* = 27,128 insertions, *n* = 23,830 deletions, and *n* = 147 inversions. Amongst all BJ (*n* = 29,258) and IMR-90 (*n* = 35,116) SVs (**Figure 6C**), *n* = 22,679 (77.5%) and *n* = 28,385 (80.8%) were heterozygous (and therefore haplotype-specific), respectively. A total of *n* = 13,269 SVs (45.4% of BJ and 37.8% of IMR-90) were shared between cell lines. Conversely, *n* = 15,989 (54.6%) of SVs found in BJ were specific to BJ whereas *n* = 21,847 (62.2%) of SVs found in IMR-90 were specific to IMR-90, suggesting that most SVs were cell-line specific. These results demonstrate that most SVs detected were heterozygous and cell-line specific.

Our analysis of these variants was based on direct inference from the assembled haplotypes. To validate the SVs using traditional alignment-based methods, we investigated three specific examples by plotting read alignments across the SV breakpoints (**Figure 6D**). We selected a 4,636 bp insertion on chromosome 2 shared by all four haplotypes, a 4,459 bp insertion on chromosome 5 present in both haplotypes of IMR-90, and a 4,342 bp deletion on chromosome 10 present in both haplotypes of BJ.

Having established the SV landscape for BJ and IMR-90, we intersected our large variant calls with the HPRC assemblies to seek additional support. After merging large insertions and deletions we found that 41% of SVs were previously reported in the *n* = 44 Year 1 HPRC samples (**Figure 6E**), indicating that many of the BJ and IMR-90 variants are also found in the pangenome. The lower proportion of large SVs found in HPRC compared to SNVs and indels found in gnomAD likely reflects the relatively smaller size of the Year 1 HPRC cohort compared to the *n* = 76,156 distinct individuals represented in the gnomAD database.

To determine which HPRC samples were most similar to BJ and IMR-90 in terms of SVs, we categorized the HPRC insertion and deletion SVs based on whether they occurred in BJ and IMR-90 (**Supplementary Figure 8A**). We found that BJ had the most shared variants with the admixed American (AMR) samples, while IMR-90 had the most shared variants with the African (AFR) samples. This prompted us to perform a genome-wide ancestry analysis of the BJ and IMR-90 assemblies (**Supplementary Figure 8B-C**). Although we found that most of both BJ and IMR-90 were assigned as European, IMR-90 contained more genomic regions than BJ that were categorized as AFR.

Intriguingly, HPRC samples showed on average *n* = 7,509 (26.58%) unique SVs that were not detected in any other samples. This contrasted with the *n* = 14,805 (58%) and *n* = 18,357 (59%) SVs that were uniquely observed in BJ and IMR-90 respectively and not in any HPRC samples. This higher variant burden potentially reflects variation acquired *in vitro*, motivating cell line assembly and underlining the importance of this work.

### Personalized reference genomes vastly improve read mapping and phasing from historical and newly generated data

To establish the benefit of using high-quality personalized assemblies relative to non-matching reference genomes, we performed traditional alignment-based sequencing analysis using outside datasets that were not used to generate the assemblies. To this end, we downloaded publicly available BJ and IMR-90 whole genome sequencing data from NCBI, ChIP-seq data from ENCODE, and generated 286 million and 138 million read pairs of RNA-seq data in-house for BJ and IMR-90 respectively (**Data Access**).

First, we reasoned that short-read data can be phased by simply aligning to the diploid genomes and using a mapping quality threshold (MAPQ ≥ 10) to keep reads that could be confidently mapped to a single haplotype. Using this method, we were able to phase 19.1% of the BJ Illumina read pairs and 25.6% of the IMR-90 Illumina read pairs. However, in regions where hap1 and hap2 are quite similar but not identical, the mapping quality will be low even if the read can be phased due to the presence of a few differences.

The alignment score (AS) takes into consideration both the length of the alignment and the edit distance between the reads and the reference. Thus, we redid the phasing analysis using the alignment score and categorized each read pair into one of five groups. A read pair was categorized as “not properly mapped” if it didn’t map to either haplotype, as “mapped to the sex chromosomes” if it mapped to chrX or chrY in either haplotype, as “homozygous” if the alignment score was the same for both haplotypes, and as “hap1” or “hap2” if the alignment score was higher for hap1 or hap2 respectively.

For the read pairs categorized as hap1 or hap2, we compared the alignment scores for concordant and discordant read pairs (**Figure 7A-B**). We found that the alignment score distributions for concordant versus discordant read pairs were significantly different. Indeed, the median of the BJ concordant read alignment scores (302) was higher than that of the discordant reads (292). Similarly, for IMR-90 the median of the concordant read alignment scores (300) was higher than that of the discordant reads (290). In BJ, 4 million more read pairs reached the maximum alignment score in the concordant alignment (**Supplementary Figure 9**). Overall, 23.2% and 27.5% of 150 bp paired-end short reads could faithfully be phased to a haplotype for BJ and IMR-90 respectively using straightforward read alignment to the matching cell line (**Supplementary Table 3**), showcasing the advantage of *de novo* diploid assembly.

**Figure 7:**
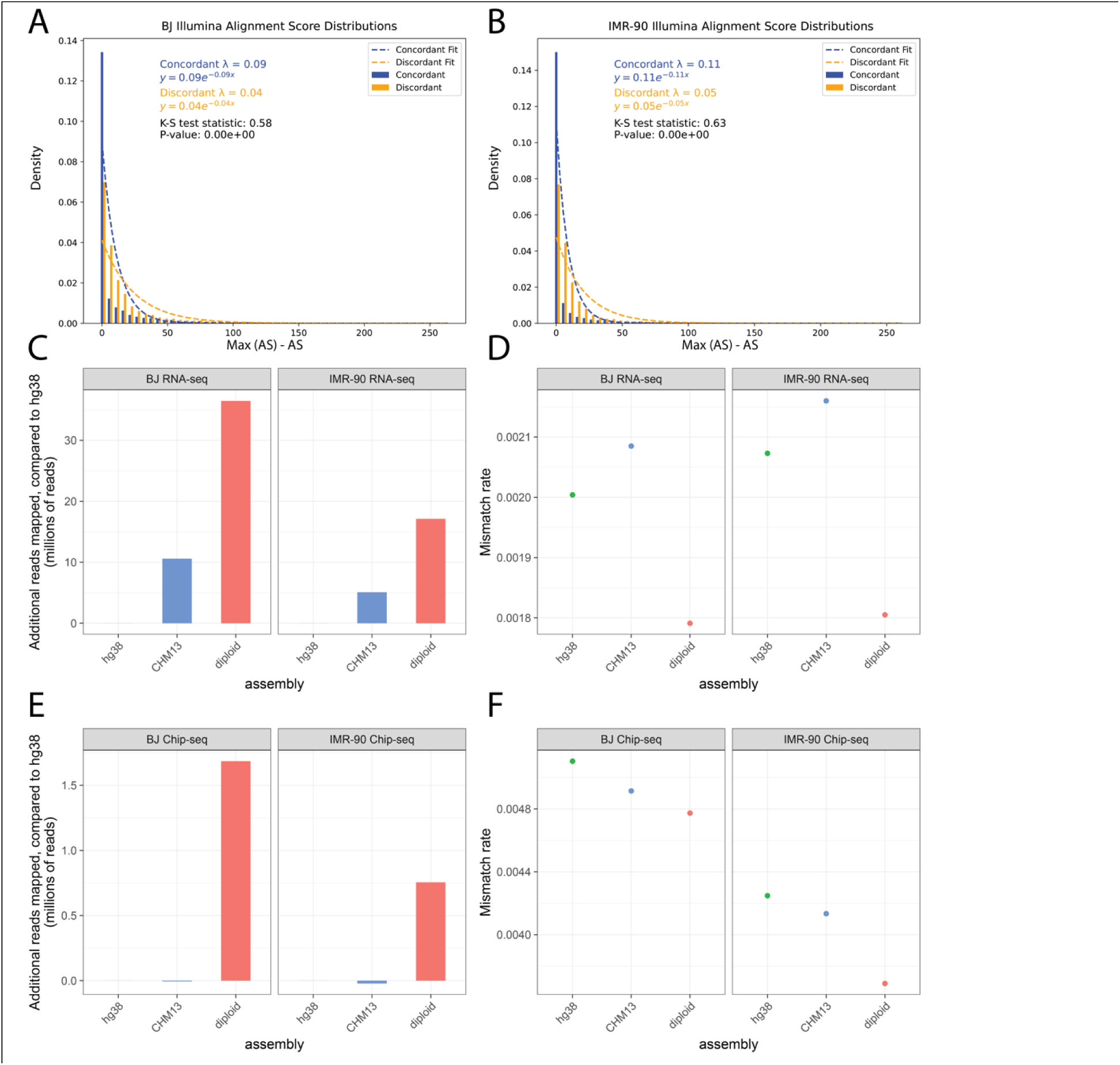
External validation of the post-curation assemblies. **A and B** Exponential distributions of the concordant and discordant read alignments of the publicly available Illumina data when mapping to the matched diploid assemblies, for BJ and IMR-90 respectively. The two distributions are statistically significantly different from each other (two-sample Kolmogorov-Smirnov tests). **C and E** Bar charts showing the number of additional mapped reads compared to the GRCh38 reference for the RNA-seq and ChIP-seq data respectively. **D and F** Dot plots showing the mismatch rate of the aligned reads, for the RNA-seq and ChIP-seq data respectively.

Next, we aligned 150 bp paired-end BJ and IMR-90 RNA-seq data to the GRCh38 reference genome, the T2T-CHM13v2.0 reference genome, and the matching *de novo* diploid assembly. After filtering for primary alignments only, we counted the differences in the number of reads mapped. Compared to the GRCh38 reference, the T2T-CHM13v2.0 reference performed slightly better, with over 10 million and 5 million more reads mapped for BJ and IMR-90, respectively (**Figure 7C**). Using the matched diploid assembly yielded even better results with over 36 million and 17 million additional reads properly mapped when compared to the GRCh38 reference for BJ and IMR-90, respectively. Comparing the subset of RNA-seq reads that mapped to all assemblies we observed the lowest mismatch rates in reads mapped to a matched assembly, suggesting that mapping reads to a personalized assembly provides better mapping and higher confidence alignment scores (**Figure 7D**).

Finally, we observed a similar trend examining alignment results of publicly available 36 bp single-end ChIP-seq data. Personalized diploid assemblies outperformed standard reference genomes in terms of number of reads aligned and mismatch rates of aligned reads (**Figure 7E-F**). In conclusion, personalized diploid assemblies can substantially improve the quality of downstream functional analyses and discoveries.

## Discussion

Our study presents the first diploid *de novo* chromosome-scale genome assemblies of the widely used human fibroblast cell lines BJ and IMR-90. These assemblies resolve complex regions of the genome in a haplotype-aware manner and reveal unique variation not found in the human reference genome or pangenome. Importantly, our assemblies facilitate improved read mapping and phasing which can yield exciting new results from the massive amount of sequencing data that has already been generated for these lines over the past several decades. Our findings underscore the importance of diploid genome assembly on widely used *in vitro* model systems.

We selected BJ and IMR-90 fibroblasts for this study because they are genetically unperturbed non-immortalized cell cultures that were directly derived from primary patient materials. Both lines harbor functional telomere maintenance mechanisms and are capable of undergoing telomere-dependent replicative senescence making them physiologically relevant models for studying telomere biology.

This key distinction contrasts our BJ and IMR-90 assemblies to ongoing efforts to establish phased and chromosome-scale genome assemblies utilizing transformed or immortalized cells in order to ensure ample amounts of input material. Generating transformed lymphoblastoid cell lines (LCLs) generally involves infecting patient-derived peripheral blood mononuclear cells (PBMCs) or isolated B cells with Epstein-Barr Virus (EBV) whereas immortalization typically requires viral transduction of human telomerase reverse transcriptase (hTERT). However, neither EBV-transformation nor hTERT-immortalization are entirely without side effects. For example, EBV-transformed cells can acquire spontaneous genetic or epigenetic changes, including chromosomal instability (e.g., chromosome gains, losses, or structural variation), telomere elongation, or changes in gene expression. Although care can be taken to minimize passaging of cells following transformation in order to reduce the risk of unintended side-effects, *in vitro* EBV-induced changes can never fully be ruled out. Our assemblies of the BJ and IMR-90 therefore present the first long-read assemblies of unperturbed human cell lines with a finite replicative potential.

A critical advance made by our study is the application of our assemblies to historical sequencing data. As a proof-of-concept, we were able to confidently phase a quarter of publicly available sequencing reads and correctly map tens of millions of otherwise unmappable reads. It is worth noting that our methods are rudimentary and future pangenome graph-alignment approaches will almost certainly prove to repurpose this data in an even more elegant fashion.

The publicly available assemblies and variant calls from this study will serve as a valuable resource for the research community, facilitating improved rigor in studies using these model systems. Future studies could focus on the functional characterization of variants that were identified in BJ and IMR-90 that have not been previously observed in the population. Perhaps cell-line specific variants in BJ are pertinent to skin fibroblasts, while those in IMR-90 are pertinent to lung fibroblasts. It is also possible that a subset of cell-specific variants occurred during *in vitro* passaging of these lines. Finally, researchers could use our diploid assemblies to study allele-specific gene regulation, for instance to investigate escape from X-chromosome inactivation in 46,XX IMR-90 cells. Our work highlights the importance of considering the cell-line specific genomic context in experimental design and interpretation.

## Methods

### Ethics Statement

A central consideration for this study was to ensure that we were free to publicly release the BJ and IMR-90 assemblies. There are ethical considerations around the public release of the data because these commercially purchased cell lines were derived before formal research use consent for biospecimens collected for clinical purposes was widely adopted. At that time, patient-derived biological materials were often used in research without explicit consent for future applications. This is likely the case for these lines, therefore WCG IRB was asked to review the proposal to share the data. The IRB assessed the risks of identifying the original subjects or their immediate family members, as well as the potential risks to groups or populations associated with submitting data to NIH-designated repositories and its subsequent sharing. Critically, openly releasing this data will significantly benefit many researchers using BJ, IMR-90, or derivative models in their laboratories, including studies of basic cell biology, disease modeling, and drug development.

Importantly, both BJ and IMR-90 are part of the National Institute of Health (NIH) Common Fund 4D Nucleome Project and the National Human Genome Research Institute (NHGRI) ENCODE Project. Large amounts of sequencing data on these cell lines are currently already publicly available through these consortia. After presenting all this information to the IRB, it was determined that the benefits of sharing the data publicly outweigh the risks and we were permitted to release our results.

### Cell Lines

BJ and IMR-90 fibroblasts were received from Michael Berens and Haiyong Han at the Translational Genomics Research Institute. They were cultured in Eagle’s Minimum Essential Medium (ATCC, Cat. No. 30-2003) + 10% FBS (VWR, Cat. No. 97068-091) according to ATCC subculturing procedures. Both cell lines were authenticated by STR profiling at ATCC (https://www.atcc.org/). Cells for sequencing were harvested with 0.25% Trypsin-EDTA (Thermo Fisher, Cat. No. 25200072), washed with 1x PBS pH 7.4 (Thermo Fisher, Cat. No. 10010023), counted on a Countess™ II FL Automated Cell Counter (Thermo Fisher, Cat. No. AMQAF1000), and used as input for NEB Monarch HMW gDNA isolation, Hi-C crosslinking, RNeasy Plus Mini isolation, or frozen as dry pellets and stored at -80 ℃ for use in Oxford Nanopore Technologies Ultra Long library preparations. Estimated cumulative population doubling level (PDL) calculations were performed using PDL = 3.32 (logXe-logXb) + S where log is the base 10 logarithm, Xb is the cell number at the beginning of the incubation period, Xe is the cell number at the end of the incubation period, and S is the starting PDL (ref ATCC Animal Cell Culture Guide ACCG-042024-v08, p6).

### gDNA Isolation & QC

BJ and IMR-90 cells were used as input for gDNA isolation using the NEB Monarch HMW DNA extraction kit for cells & blood (New England Biolabs, Cat. No. T3050L) following manufacturer’s instructions. HMW DNA was quantified by Nanodrop and Qubit^TM^ 1xdsDNA High Sensitivity kit (Thermo Fisher, Cat. No. Q33231). Rough size distribution and DNA Integrity Number (DIN) were measured with Genomic DNA ScreenTape and reagents (Agilent, Cat. No. 5067-5365 and 5067-5366) using the Agilent 4150 TapeStation System (Agilent, Cat. No. G2992AA).

### PacBio HiFi Library Preparation and Sequencing

2-5 ug of HMW gDNA was used as input for SMRTbell Prep Kit V3.0 (PacBio, Cat. No. 102-182-700). Mechanical shearing of gDNA was performed with Megaruptor 3 (Diagenode, Cat. No. B06010003) using the standard protocol with target size of 15-20 kb. Fragmentation was analyzed by Femto Pulse System (Agilent, Cat. No. M5330AA). SMRTbell libraries were sequenced on the PacBio Revio platform with SMRT Cell tray (PacBio, Cat. No. 102-202-200), loaded at 240-260 pM, with 30-hour movie collection times. For BJ, 2 libraries were made and run on 3 flowcells to reach >30x coverage of the genome. For IMR-90, 2 libraries were made and run on 2 flowcells to reach >30x coverage of the genome.

### Hi-C Crosslinking

1-3 x 10^6^ freshly harvested cells were crosslinked using 1% methanol-free formaldehyde (Pierce, Cat. No. 28906) in 1x PBS pH 7.4 for 10 min with gentle agitation on a platform rocker. Crosslinking reactions were quenched with 200 mM Glycine at RT 5 min with rotation at 10 RPM followed by 15 min incubation on ice. Crosslinked cells were washed in 1x PBS pH 7.4, frozen as dry pellets, and stored at -80 ℃.

### Hi-C Library Preparation and Sequencing

Hi-C assays were performed on frozen crosslinked cell pellets using the Arima High Coverage HiC Kit (Arima Genomics, Cat. No. A101030) and libraries were prepared using Arima Library Prep Kit v2 (Arima Genomics, Cat. No. A303011) per manufacturer’s instructions. 1.5 µg of proximally-ligated DNA suspended in 130 µL of the elution buffer was added to a microTUBE (Covaris, Cat. No. 520045) and sonicated using a Covaris E220 with the following parameters: Temperature (4-7 °C), Peak Incident Power (105), Duty Factor (5%), Cycles per Burst (200), Treatment time (75 s). Post-sonication QC was performed on a TapeStation HSD5000 (Agilent Technologies, Cat. No. 5067-5592) to verify that fragment sizes were between 550-600 bp prior to size selection. Size selection was performed with AMPure XP beads (Beckman Coulter, Cat. No. A63881) to collect fragments >400 bp. 200 ng of size-selected DNA was put into biotin enrichment. The enriched products were end-repaired, adapter-ligated, and tagged with manufacturer supplied P7 and P5 8 bp index pairs using 5 cycles of PCR. The final bead-purified libraries were quantified by Qubit dsDNA HS (Thermo Fisher, Cat. No. Q33231) and TapeStation HSD5000 screentape, pooled, and re-quantified. The pool was sequenced on a NovaSeqX 10B flow cell (Illumina, Cat. No. 20085594) for 151 x 151 cycles with a 1% PhiX spike-in to 25x coverage.

### Oxford Nanopore Technologies (ONT) Ultra-Long Sequencing

Dry cell pellets of 6 x 10^6^ of each cell type were used as input for ONT-UL library preparation per Ultra-Long DNA Sequencing Kit V14 (Oxford Nanopore Technologies, Cat. No. SQK-ULK114). Libraries were loaded on PromethION Flow Cells (R10.4.1) (Oxford Nanopore Technologies, Cat. No. FLO-PRO114M). BJ and IMR-90 libraries were loaded onto 7 and 9 flowcells, respectively, to reach >20x coverage of the genome for each cell line.

### RNA Isolation & QC

For each cell pellet, total RNA was isolated with RNeasy Plus Mini (Qiagen, Cat. No. 74134) using QIAshredder columns (Qiagen, Cat. No. 79654) for homogenization, then purified with Zymo RNA Clean & Concentrator-5 (Zymo Research, Cat. No. R1015). Isolated RNA was measured for quantity with Quant-iT Ribogreen RNA Assay (Thermo Fisher, Cat. No. R11490) and quality with High Sensitivity RNA ScreenTape and buffer (Agilent, Cat. No. 5067-5579 & 5067-5580).

### Whole Transcriptome Library Preparation & Sequencing

For each RNA sample, a uniquely dual-indexed, Illumina-compatible, double-stranded cDNA whole transcriptome library was synthesized from 1 µg of total RNA with Takara Bio’s SMARTer Stranded Total RNA Sample Prep Kit - HI Mammalian kit (Takara Bio, Cat. No. 634874) and SMARTer RNA Unique Dual Index kit (Takara Bio, Cat. No. 634452). Briefly, this library preparation included ribosomal depletion, RNA fragmentation (94 °C for 3 min), and a 12-cycle indexing and enrichment PCR. Each library was measured for size with Agilent’s High Sensitivity D1000 ScreenTape and buffer (Agilent, Cat. No. 5067-5584 & 5067-5603). 1 uL of each library was combined into a non-equimolar pool which was then measured for size with TapeStation and concentration with Roche’s KAPA SYBR FAST Universal qPCR Kit (Roche, Cat. No. KK4824), diluted to 70 pM, then loaded into an iSeq flow cell cartridge (Illumina, Cat. No. 20031371) with a 1% v/v PhiX Control v3 spike-in (Illumina, Cat. No. FC-110-3001), and sequenced to 101 x 9 x 9 x 101 cycles. Passing filter cluster counts per library from these iSeq data were used to make a re-balanced pool which was subsequently measured for size and concentration, diluted to 750 pM with a 1% v/v PhiX Control v3 spike-in, denatured and further diluted, loaded into a NovaSeq X flow cell lane (Illumina, Cat. No. 20085594), and sequenced at 151 x 8 x 8 x 101 cycles with a final flow cell concentration of 150 pM. Libraries were sequenced to at least 50 M read pairs (or 100 M paired-end reads). The low diversity of the first three bases in Read 1 of these libraries was balanced in both the iSeq and NovaSeq X pools with other library types that had a higher diversity across those bases; otherwise, a 20% PhiX spike-in would have been needed for both pools.

### Multi-color Metaphase Fluorescence In Situ Hybridization (mFISH)

BJ and IMR-90 cells were cultured to 70% confluency, harvested with 0.5% Trypsin-EDTA (Gibco, Cat. No. 15400054), collected by centrifugation at 1,500 rpm for 5 minutes, and resuspended with 0.075 M KCl (Invitrogen, Cat. No. AM9640G) for 10 minutes. Ice-cold Methanol: Acetic Acid solution was used to resuspend and fix the cells. 30 µL of cell suspension was dropped onto pre-cleaned slides and hybridized to whole chromosome PAINT probes (MetaSystems, Cat. No. D-0125-060-DI) for 48 hours, according to the manufacturer’s protocol. Coverslips were mounted on slides with ProLong Glass Antifade with NucBlue (Thermo Fisher, Cat. No. P36981). Images were acquired using a Zeiss AxioImager Z2 equipped with the MetaSystems *Metafer* Slide Scanning Platform (MetaSystems, *Metafer5*).

### Karyogram Analysis

Karyograms were generated with the Interactive Karyotyping system (Ikaros) module, a feature of the Metafer system. Captured images were processed by background subtraction, object thresholding, object separation, and pixel calling. False colors were assigned to each chromosome for clear identification. Chromosome assignments were confirmed by comparing fluorescent spectra to the manufacturer key and matching each chromosome to its respective ideogram. Reports for each cell line were generated and exported from the software.

### Sequencing QC

HiFi read ubam files were converted to fasta using Samtools (version 1.21)^34^. ONT-UL reads were basecalled using Dorado (version 0.7.2+9ac85c6)^35^ with the dna_r10.4.1_e8.2_400bps_sup@v5.0.0 model to produce duplex ubam files. The ubams were converted to fastq using Samtools (version 1.21)^34^. We retained the reads with the dx tag set to 1 or 0 to keep the duplex reads and the simplex reads without duplex offsprings. The ONT-UL reads were error corrected using Dorado (version 0.7.2+9ac85c6)^35^. The first five bases from the 5’ end of both read 1 and read 2 of the Hi-C data were trimmed using cutadapt (version 4.5)^36^.

We constructed k-mer histograms for the long-read sequencing data using kmc (version 3.2.4)^37^. We analyzed the k-mer histograms using GenomeScope (version 2.0.1)^19^ to estimate the coverage, heterozygosity, and error rate of the data. We used NanoPlot (version 1.44.0)^18^ to generate read length vs average read quality KDE plots and get the total sequencing coverage, read length N50, and median quality scores of the sequencing data. We generated coverage versus read length plots using a python script. We aligned the Hi-C data to the T2T-CHM13v2.0 reference^12^ with bowtie2 (version 2.5.4)^38^ and used hic_qc (version 1.0)^20^ to plot the distribution of long range contacts.

### Genome Assembly

We generated phased diploid assemblies of the cell lines using Verkko (version 2.2.1)^21^. Prior to assembly, the ONT-UL data was error corrected using Dorado (version 0.8.1+c3a2952)^35^. The HiFi reads and the error corrected ONT-UL reads were given as hifi inputs to Verkko. The ONT-UL reads and the Hi-C reads were given as nanopore and hic inputs respectively to Verkko.

### Assembly Curation

Hi-C reads were aligned to the pre-curation diploid assemblies (including hap1, hap2, and the unassigned reads) and processed according to the Arima Mapping pipeline (A160156 v03)^39^. In summary, Hi-C reads were aligned to the pre-curation diploid assemblies using BWA mem with default parameters (version 0.7.18-r1243-dirty)^40^ . The bam files were filtered using the filter_five_end.pl script to retain only the portion of chimeric reads that map in the 5’-orientation. The two_read_bam_combiner.pl script was used to combine the single-end Hi-C reads. Picard (version 3.1.1)^41^ was used to add read groups to the bam files and remove optical duplicates. Hi-C contact maps were created with PretextMap (version 0.1.9)^42^.

Using PretextView (version 0.2.5)^43^ for visualization of this contact map, we were able to detect misjoins and consolidate segments from the same chromosomes into a single scaffold. Based on this refined structure, we proceeded to generate an AGP (A Golden Path) file to define the scaffold organization. This file was subsequently used as input for the Rapid Curation 2.0 pipeline^44^ to separate the two haplotypes^45^.

### Assembly QC

Bandage (version 0.8.1)^46^ was used to view the assembly graphs from Verkko. gfastats (version 1.3.9)^47^ was used to calculate the genome size, total gap length, number of gaps, contig and scaffold NG50, and GC content of the assemblies. The NG50 (respectively LG50) is the length (number) of the shortest sequence needed to cover at least 50% of the total reference assembly. The NGx and LGx curves plot the entire range of NG and LG values. The auN value represents the area under the NGx curve. The NGx and LGx charts were plotted using a custom R script. The auN values of the assembly were calculated using a custom Python script. The RepeatMasker (version 4.1.6)^24^ was used to profile the repeat sequences in the assemblies using the Dfam withRBRM (version 3.8) database^48^ and the ABBlast search engine.

Compleasm (version 0.2.6)^22^ with the primates lineage was used to evaluate the genome completeness by searching the assemblies for near-universal single-copy ortholog gene sequences^49^. Meryl (version 1.4.1)^23^ was used to count the 21-mers from publicly available Illumina data (see Data Availability)^50,51^. The quality of the assemblies was evaluated using Merqury (version 1.3)^23^.

The HiFi reads were aligned to the diploid assemblies using minimap2 (version 2.28-r1209)^52^ and HMM-Flagger (version 1.0.0)^13^ was used to detect any misassemblies. Liftoff (version 1.6.3)^53^ was used to lift the annotation of the T2T-CHM13v2.0 reference^12^ separately to each haplotype of the assemblies.

The computational karyograms were generated by coloring the 31-mers of the assemblies based on which chromosome arm of T2T-CHM13v2.0 they are located. Each pixel represents 250 Kbp of sequence and is colored according to majority rule. Centromeric 31-mers are colored orange, novel 31-mers not present in T2T-CHM13v2.0 are colored white, 31-mers that occur on multiple chromosome arms are colored grey, and telomeres are notated with blue circles on the chromosome ends. Each chromosome has a unique color, with the p-arm and q-arm of a given chromosome colored light and dark respectively. The sequences are clustered into their corresponding chromosomes and haplotypes. Sequences smaller than 1 Mbp are filtered out.

ntSynt (version 1.0.2)^28^ was used to produce the multi-sample synteny plot between the T2T-CHM13v2.0 reference and the four haplotypes of the assemblies. SVbyEye (version 0.99.0)^30^ was used to produce the synteny plots of hap1 versus hap2 for BJ and IMR-90. The synteny plots of the IMR-90 haplotypes to the T2T-CHM13v2.0 reference showing the chromosome 8 inversion were produced by SyRI (version 1.6.3)^31^.

SyRI (version 1.6.3) was used to call variants of each of the four assembly haplotypes against the T2T-CHM13v2.0 reference. SNVs and indels were compared against the gnomAD^32^ dataset. The resulting intersection data were reformatted into a binary matrix for UpSet plot analysis using ComplexUpset (version 1.3.3)^54^. Variants exclusive to gnomAD were excluded from the UpSet plot. The SVs across the haplotypes were merged with Jasmine (version 1.1.5)^33^. SVs with fewer than 50 bp were removed. The stacked upset plot of SV counts was generated using the UpSetPlot Python package (version 0.9.0)^55,56^. Individual SVs were visualized using IGV (version 2.18.4)^57^ and Ribbon (version 2.0)^58^.

For the ancestry analysis, RFMix2 tool^59^ as used to infer local ancestry for each haplotype of the assemblies. As a reference panel, the phased VCF from the 1000 Genomes Project^60^ were downloaded. The target dataset consisted of phased variants generated using HiPhase^61^ and DeepVariant^62^. Local ancestry inference was performed using RFMix2 with default parameters. To visualize chromosome-level ancestry, the AncestryGrapher toolkit^63^ was employed, which provides chromosome painting representations.

fastp (version 0.24.0)^64^ was used for adapter trimming, deduplication, and overrepresented sequence analysis of the ChIP-seq and RNA-seq data. BWA (version 0.7.18-r1243-dirty)^40^ was used to align the ChIP-seq reads to the GRCh38^11^ and T2T-CHM13v2.0^12^ references and to the post-curation diploid assemblies. HISAT2 (version 2.2.1)^65^ was used to align the RNA-seq reads to the GRCh38 and T2T-CHM13v2.0 references and to the post-curation diploid assemblies. Picard (version 3.1.1)^41^ was used to mark optical duplicates. GATK (version 4.5.0.0)^66^ CollectAlignmentSummaryMetrics was used to determine the number of mapped reads (PF_READS_ALIGNED) and the mismatch rate (PF_MISMATCH_RATE).

BWA (version 0.7.18-r1243-dirty)^40^ was used to align the Illumina reads separately to each post-curation haplotype. The reads were phased based on which alignment (hap1 or hap2) had a higher alignment score (AS tag) using a custom Python script. The reads were categorized as not properly mapped if they didn’t map to either haplotype, as mapped to the sex chromosomes if they mapped to chrX or chrY in either haplotype, as homozygous if the alignment score was the same for both haplotypes, as hap1 if the alignment score was higher for hap1, and as hap2 if the alignment score was higher for hap2.

For those reads that could be phased, the distribution of AS scores was plotted as a histogram. Specifically, we created a concordant distribution to include all the AS scores for the concordant alignments (hap1 reads aligned to hap1 or hap2 reads aligned to hap2) and a discordant distribution to include all the AS scores for the discordant alignments. For each distribution, we fit the best exponential distribution using the statistical functions in SciPy (version 1.10.1)^67^ in Python 3.11.3. We also used a two-sample Kolmogorov-Smirnov test^68^ to determine whether the distributions came from the same distribution.

Centromere annotation was done by intersecting the outputs derived from RepeatMasker^24^ and HumAS-HMMER for AnVIL^69^ to identify the position of the centromeres and the Higher-Order Repeats (HORs) and Superfamilies (SF) within each centromere. RepeatMasker was run with default parameters, while HumAS-HMMER for AnVIL was applied as described in Altemose et al., 2022^70^. To generate heatmaps showing the variation between centromeres, StainedGlass (version 6.7.0)^27^ was run with the following parameters: window=5000 and mm_f=30000. Monomer-by-monomer annotation and structural variants detection within the live HORs was done using StV^70^.

The Foreign Contamination Screening (FCS) tool (version 0.5.4)^71^ from NCBI was used to detect adaptor and vector contamination in the post-curation assemblies. FCS was also used to screen for non-human sequences in the post-curation assemblies.

## Supplementary Figures

**Supplementary Figure 1:**
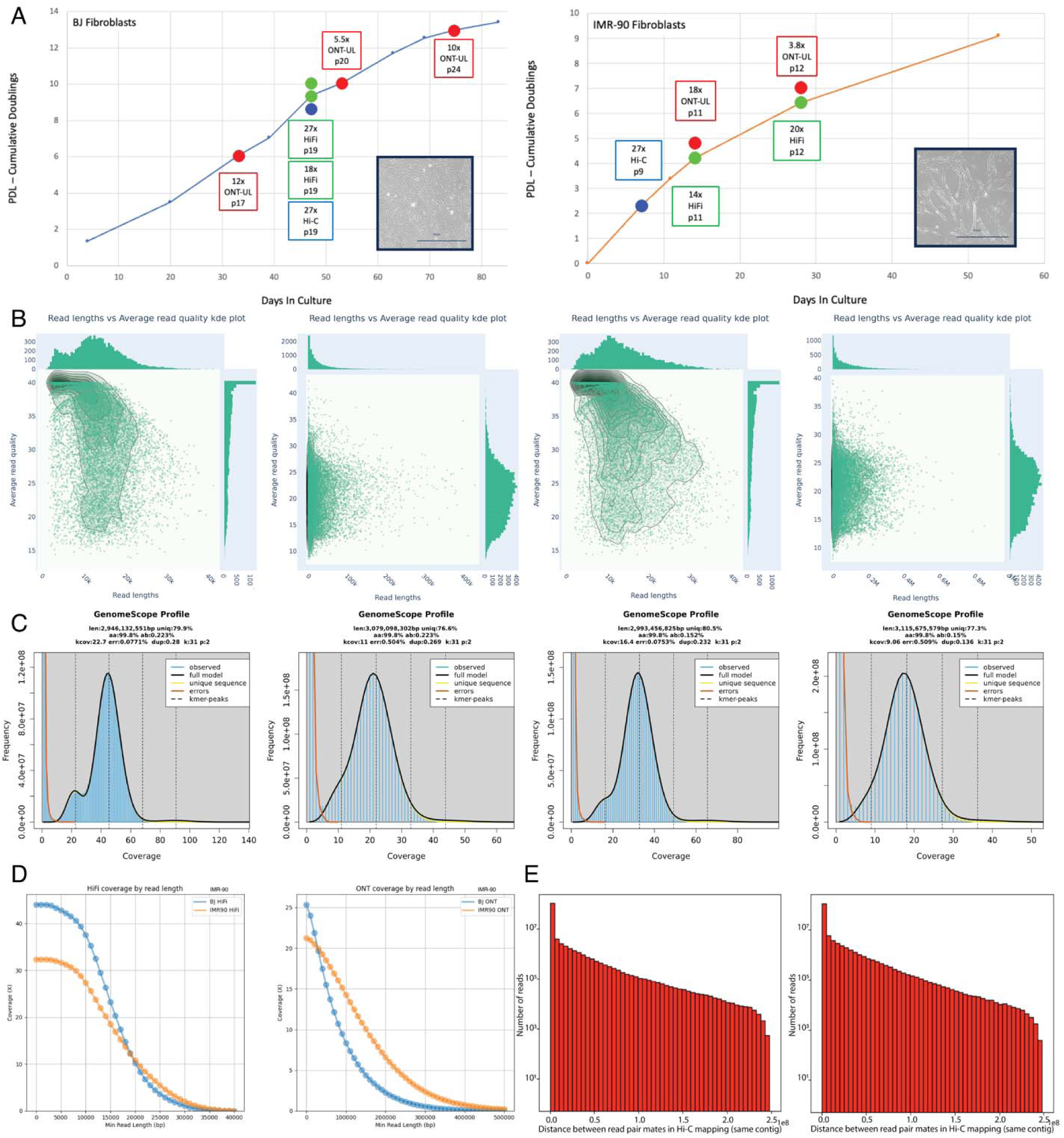
A BJ and IMR-90 cell culture growth curves. Plots show cumulative doublings of cultured fibroblasts with time points where cells were harvested for different sequencing platforms (HiFi, Hi-C, ONT-UL, and RNA-seq). Passage number and sequencing coverage for each harvested time point are shown in boxes at each respective time point. Microscope images with representative examples of each cell type are shown in inset of each growth curve. **B** NanoPlot read length vs average read quality KDE plots for BJ HiFi, BJ ONT-UL, IMR-90 HiFi, IMR-90 ONT-UL data respectively. **C** GenomeScope plots for BJ HiFi, BJ ONT-UL, IMR-90 HiFi, and IMR-90 ONT-UL data respectively show estimated coverage and heterozygosity. **D** Coverage vs read length for BJ and IMR-90 for HiFi and ONT-UL data, respectively. **E** Hi-C QC results for BJ and IMR-90. Growth curves and sequence data QC

**Supplementary Figure 2:**
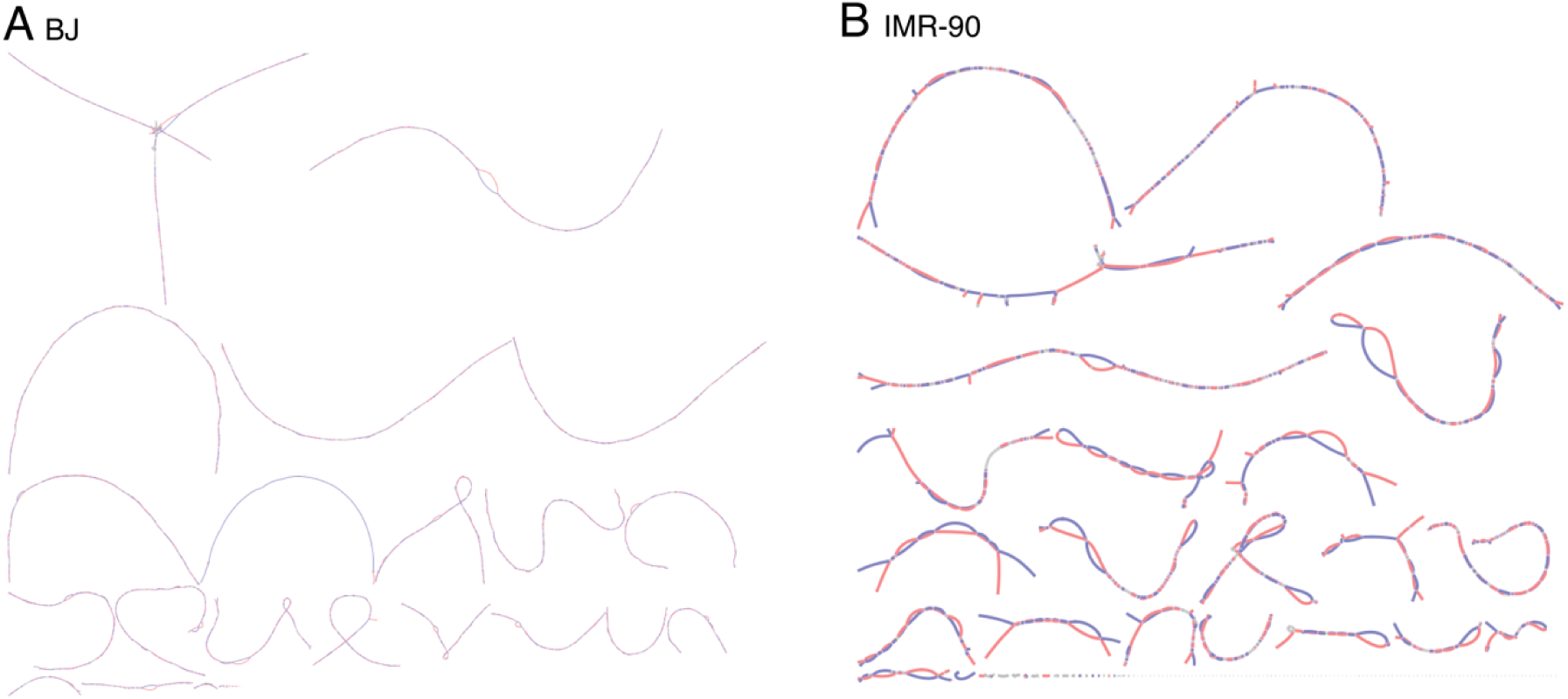
Bandage plots of the Verkko assemblies for **A** BJ and **B** IMR-90. Most of the chromosomes are in a single contiguous piece. The acrocentric chromosomes also belong to the same piece as expected. Assembly bandage plots

**Supplementary Figure 3:**
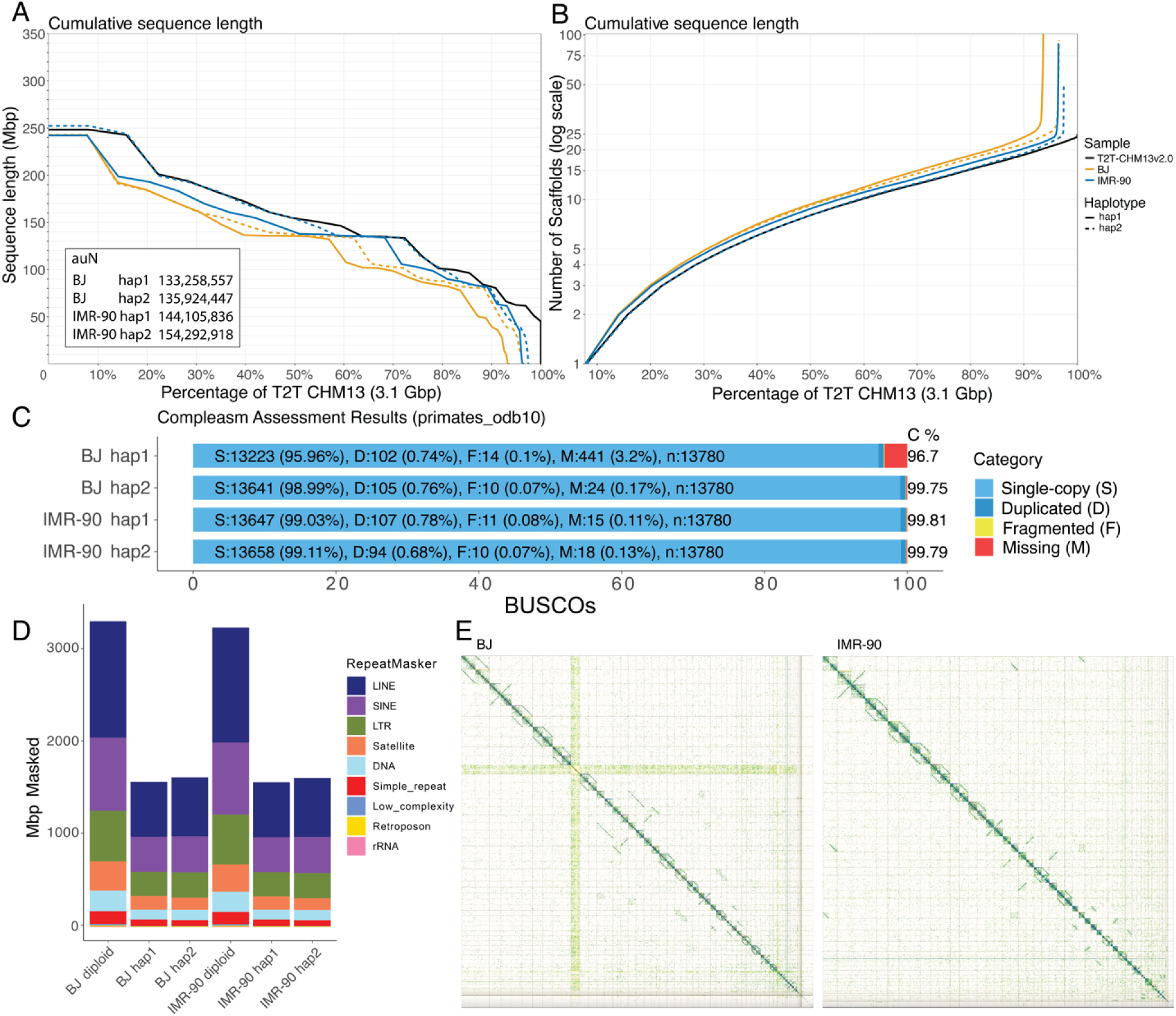
Pre-curation assembly QC. **A** NGx chart showing the contiguity of the four pre-curation assembly haplotypes (based on the genome size of the T2T-CHM13v2.0 reference). The NG50 value (when x=50%) is the length of the shortest sequence needed to cover at least 50% of the total reference assembly. In black is the NGx plot for the T2T-CHM13v2.0 reference. Overlaid on the chart are the auN metrics for the pre-curation haplotypes. **B** LGx charts showing the number of contigs needed to reach a given percentage of the human genome. **C** The compleasm results showing the estimated number of single-copy (S), duplicated (D), fragmented (F), and missing (M) near-universal single-copy ortholog gene sequences in the pre-curation haplotypes. **D** RepeatMasker results showing the number of bases in the pre-curation haplotypes that are categorized into each repeat family. **E** Hi-C maps after mapping the Hi-C data to the pre-curation assemblies. Pre-curation assembly QC

**Supplementary Figure 4:**
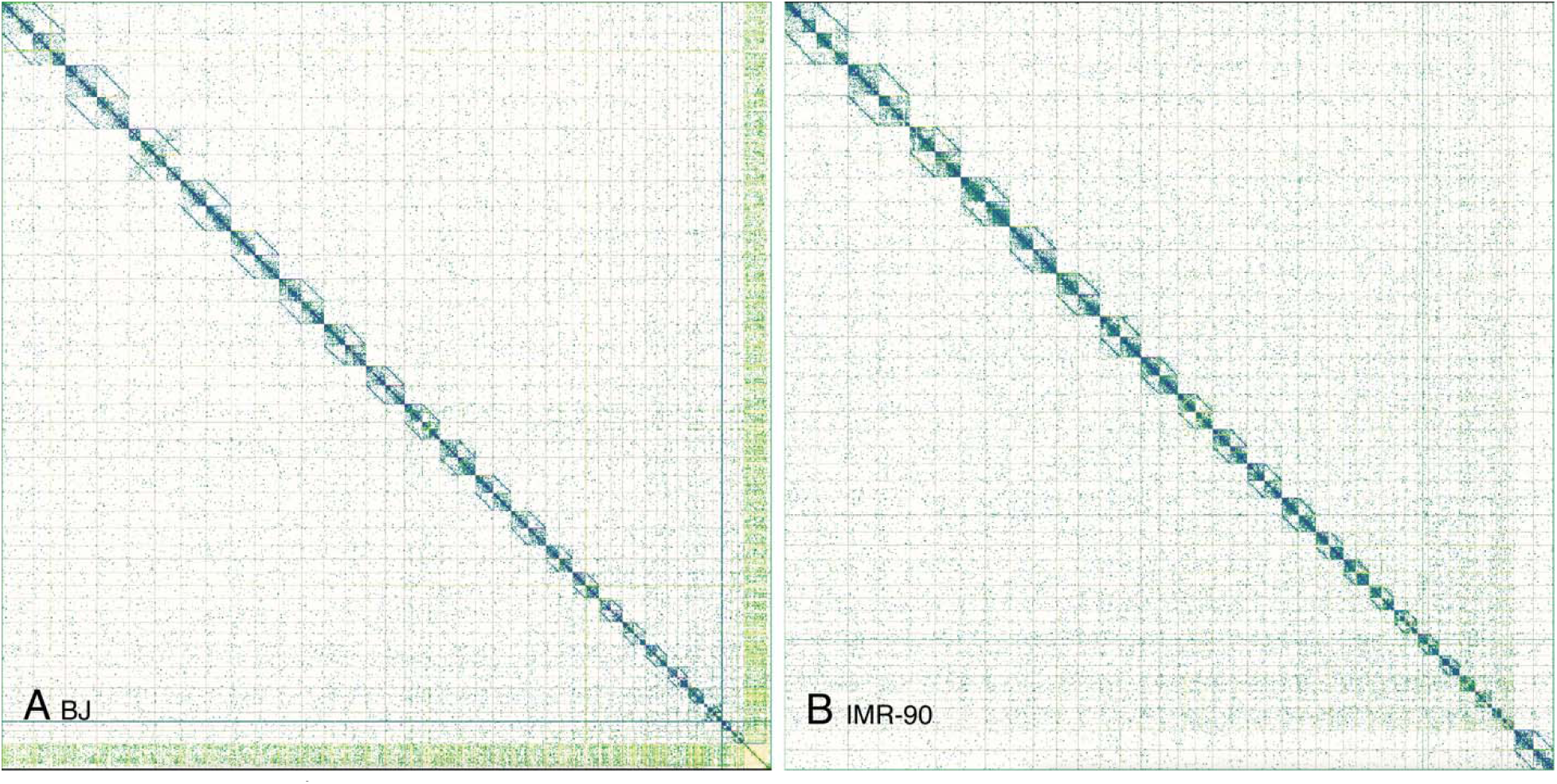
Post-curation Hi-C maps for **A** BJ and **B** IMR-90. These Hi-C maps show the characteristic diamond shape for each diploid chromosome. There are no significant off diagonal contacts. Post-curation Hi-C maps

**Supplementary Figure 5:**
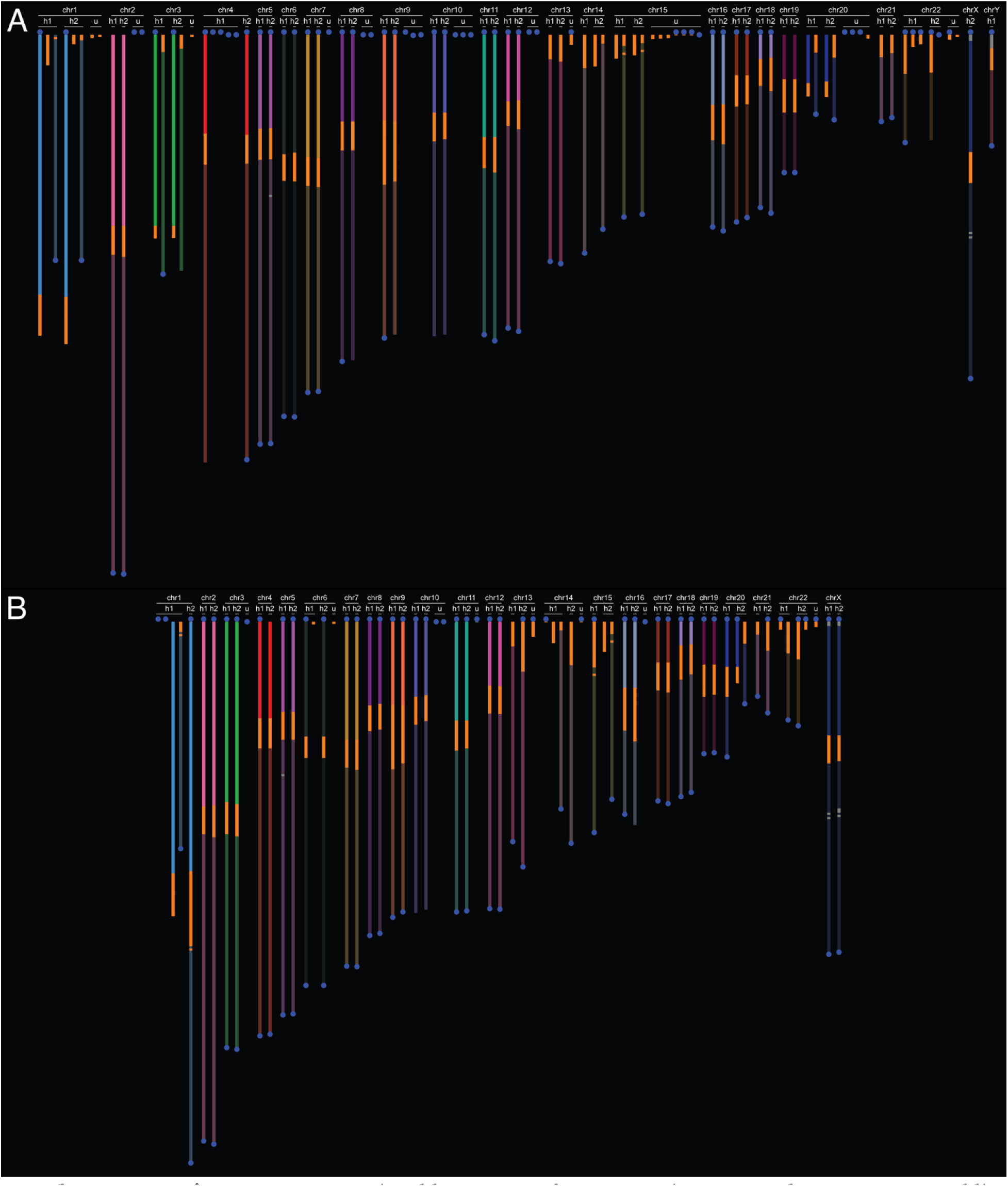
Computational karyograms for pre-curation **A** BJ and **B** IMR-90 assemblies (see **Methods**). Most of the chromosomes consist of a single contig. A few of the chromosomes consist of two contigs broken at the centromere. The sex chromosomes are assembled in a single contig with telomeres at each end. Pre-curation computational karyogram

**Supplementary Figure 6:**
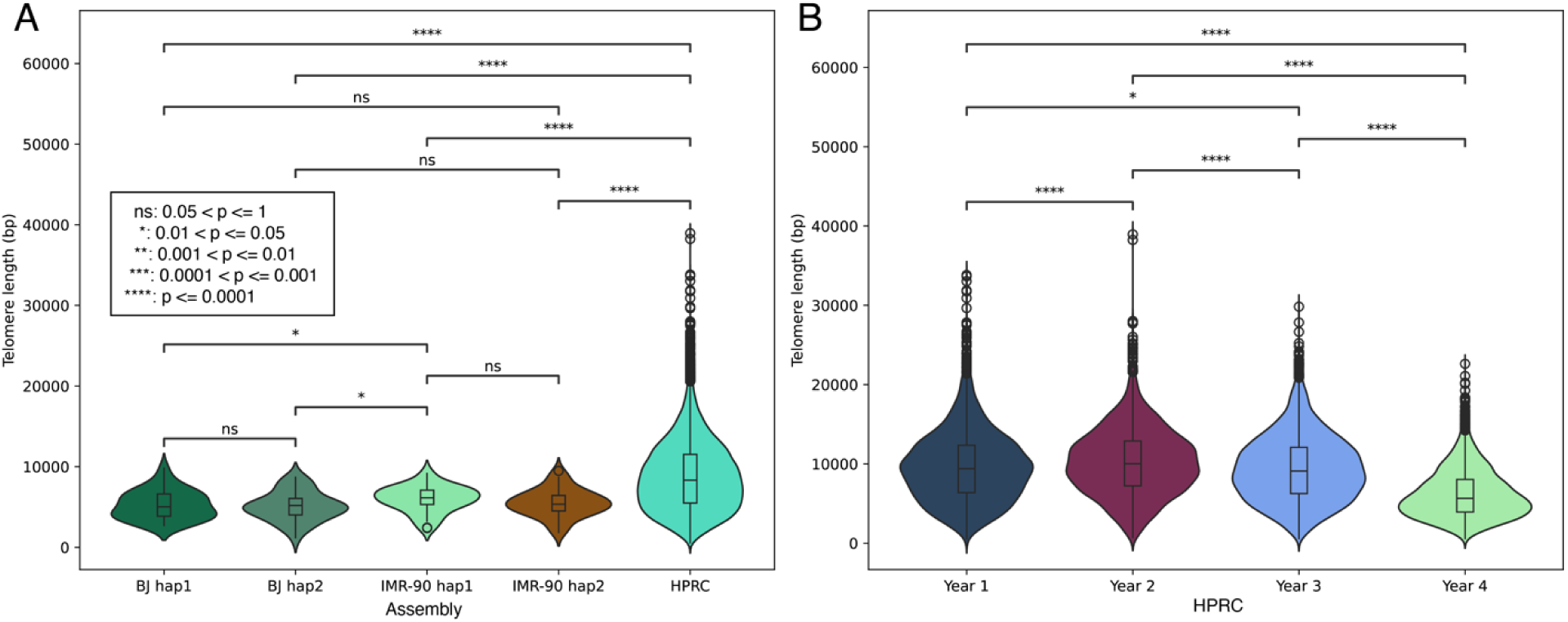
A. BJ and IMR-90 telomere lengths compared to the HPRC human pangenome. Mann-Whitney-Wilcoxon two-sided tests were performed to compare the median telomere lengths. **B** Telomere lengths in years 1, 2, 3 and 4 of the HPRC pangenome. BJ and IMR-90 telomere lengths compared to the human pangenome

**Supplementary Figure 7:**
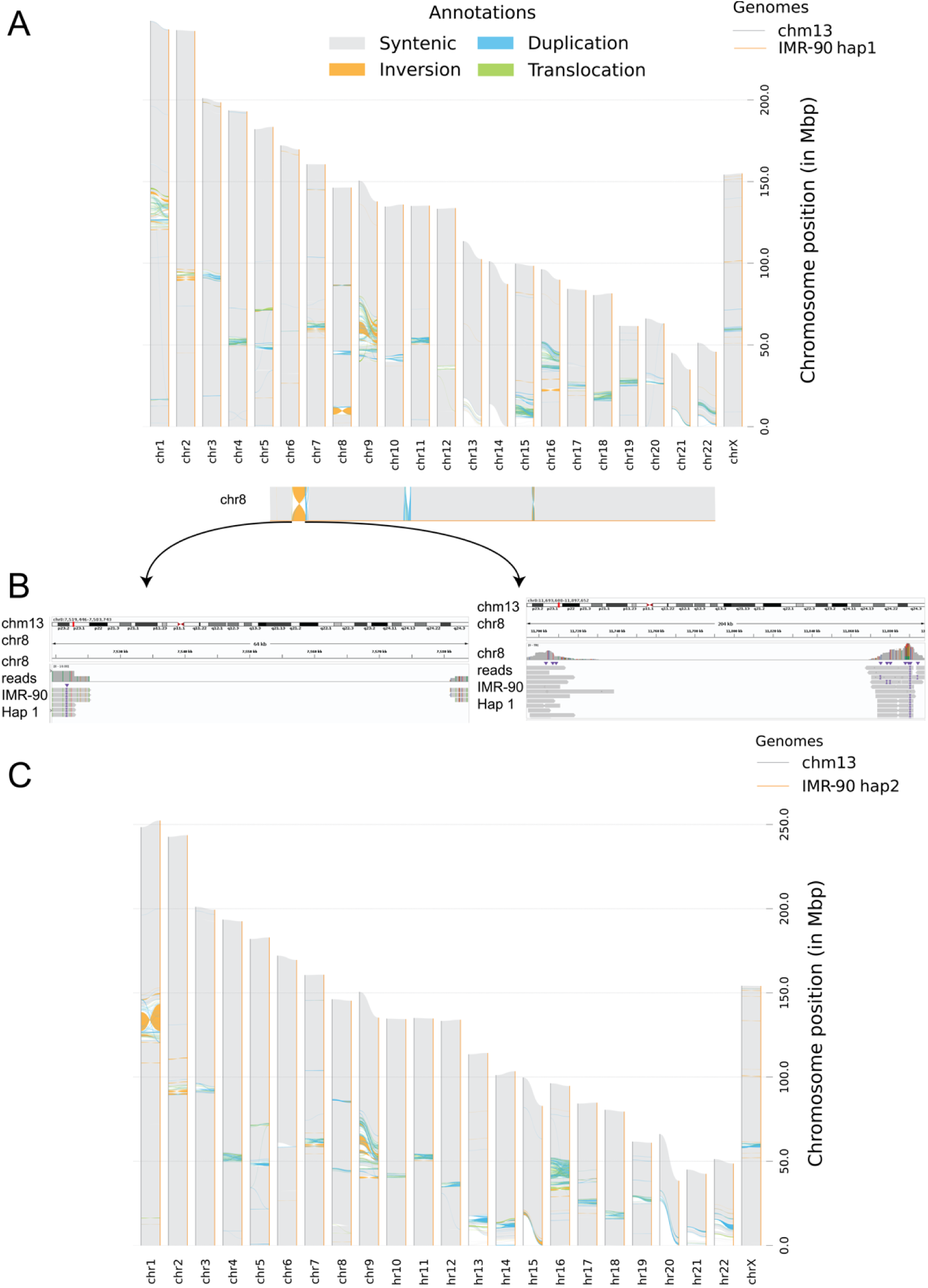
Example haplotype-specific inversion on chromosome 8 in IMR-90. **A** SyRI synteny plot of IMR-90 haplotype1 against the T2T-CHM13v2.0 reference. Inversions are in orange color. At the bottom is the zoomed-in view of the example inversion. **B** IGV screenshots of IMR-90 reads aligned to T2T-CHM13v2.0 chromosome 8 at both breakpoints of the inversion. **C** SyRI synteny plot of IMR-90 haplotype2 against the T2T-CHM13v2.0 reference showing no inversion in chromosome 8. Example haplotype-specific inversion on chr8 in IMR-90

**Supplementary Figure 8:**
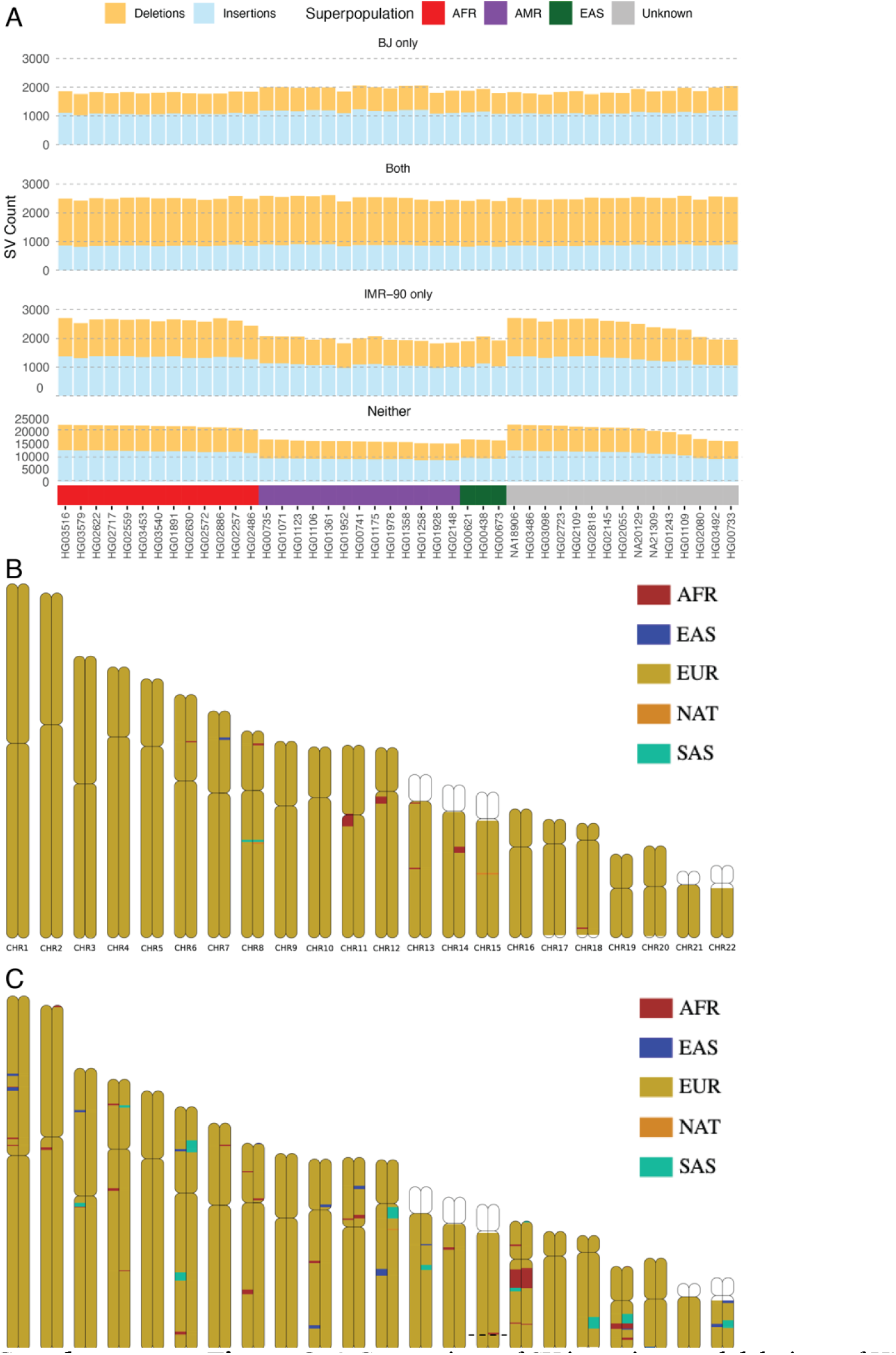
A. Comparison of SV insertions and deletions of HPRC pangenome to BJ and IMR-90. The pangenome samples are ordered by superpopulation. Pangenome variants are divided into four groups: “Both” if they are also present in both BJ and IMR-90, “BJ only” or “IMR-90” only if they are also present in that cell line but not the other, and “Neither” if they are not present in either cell line. The number of “BJ only” variants is highest for the AMR pangenome samples, while the number of “IMR-90” only variants is highest for AFR. **B** BJ chromosome painting based on ancestry. **C** IMR-90 chromosome painting based on ancestry. Comparison of pangenome and cell line SVs and ancestry analysis

**Supplementary Figure 9:**
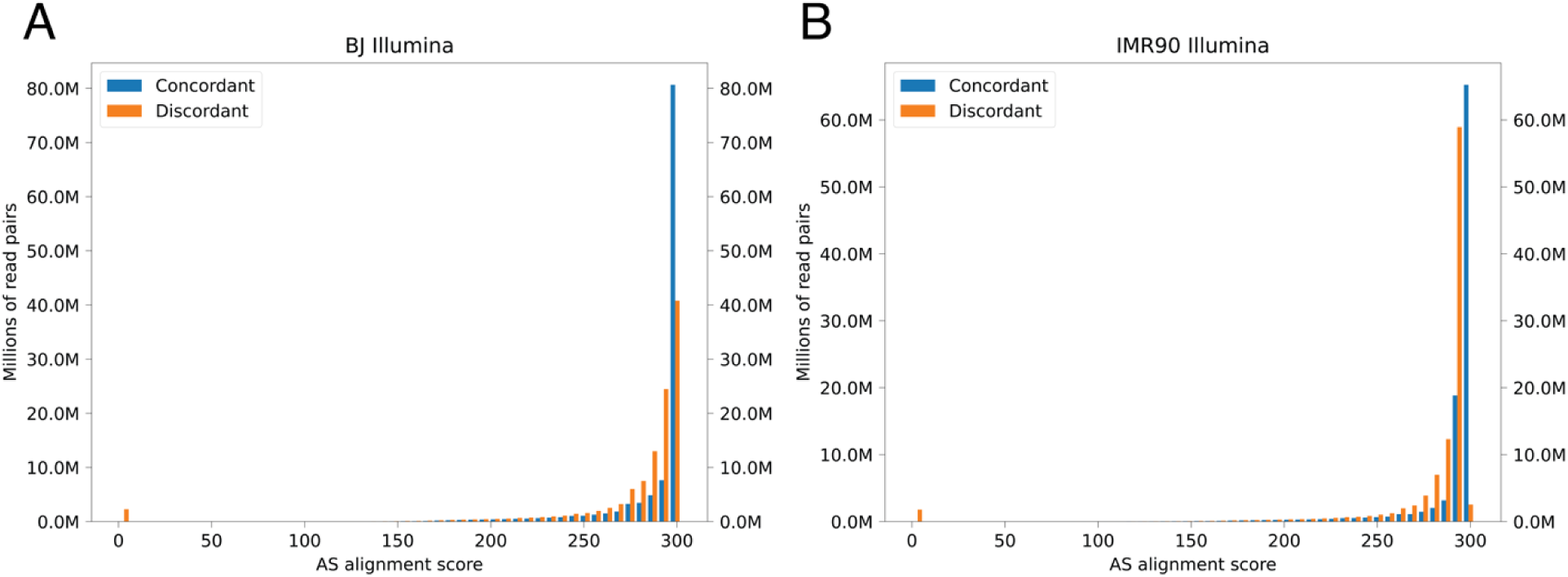
BWA alignment score (AS) distributions of concordant versus discordant alignments of Illumina reads against the post-curation **A** BJ and **B** IMR-90 assemblies. The alignment scores range from 0 to 300 for each read pair. An alignment score of 300 means that all 300 bp of the read pair match the reference exactly. Any deletions, insertions, or mismatches lower the scores. Read pairs that are not properly mapped have an alignment score of 0. These figures show that the concordant alignments (when mapping to the correct haplotype) have much higher alignment scores than the discordant alignments. Alignment score distributions

## Supplementary Tables

See Excel file.

## Data Access

The HiFi, ONT-UL, Hi-C, and RNA-seq data generated in this study have been submitted to the NCBI BioProject database (https://www.ncbi.nlm.nih.gov/bioproject/) under accession number PRJNA1214805. Publicly available Illumina data for BJ and IMR-90 was downloaded from the NCBI SRA database (BJ BioSamples ID: SAMD00260498, BioProject ID: PRJDB6632, Run ID: DRR258666; IMR-90 BioSamples ID: SAMN35312059, BioProject ID: PRJNA975175, Run ID: SRR24709136). Publicly available ChIP-seq data for BJ and IMR-90 was downloaded from ENCODE (BJ Experiment: ENCSR000DQH, Files: ENCFF001ERS and ENCFF001ESA; IMR-90 Experiment: ENCSR087PFU, Files: ENCFF591EQV and ENCFF742UXQ).

## Author Contributions

T.R.R-B. developed computational pipelines and led data analysis. Y.H. co-led data analysis and produced the final figures. E.V. performed the manual curation of the assemblies. M.J. supported data analysis. N.F. and M.M. performed cell culture, sample preparation and karyotyping. D.M.P., R.R., J.M., D.M., and S.W. performed library preparation and sequencing. T.R.R-B., Y.H., M.J.M., M.K., B.P., S.G and F.P.B. interpreted the results. K.K and S.B. oversaw research compliance. T.R.R-B., Y.H., N.F., E.V. and F.P.B wrote the manuscript. F.P.B. conceived and supervised the project. All authors reviewed, commented on and approved the final paper.

## Acknowledgements

This research includes work performed in TGen’s Collaborative Sequencing Center, a City of Hope Comprehensive Cancer Center supported shared resource (NCI-P30CA033572). This work was supported in part by funding from The Lennar Foundation. We are grateful to Drs. Michael Berens and Haiyong Han at the Translational Genomics Research Institute who provided the BJ and IMR-90 fibroblasts. We would like to thank Drs. Karen Miga, Adam Phillippy, and Justin Zook for the invaluable discussions surrounding the ethical, legal and social issues of genome assembly and data sharing. S.G. receives salary support from the Rita Levi Montalcini Award, E.V. is supported by a PON PNRR Ph.D. Fellowship from the Italian Ministry of University and Research (MUR), and work in the Giunta lab is supported by AIRC Start-Up grant 202 ID # 25189, Sapienza Excellence grant and ERC CENTROFUN grant # 101078838 to S.G.

